# Nonequilibrium States Promote One-Pot Nonenzymatic Carbon Fixation in the Reverse Tricarboxylic Acid Cycle and Amino Acid Synthesis

**DOI:** 10.64898/2026.04.29.721051

**Authors:** Yu-Hsi Lin, Jian-Hau Peng, Shao-Yu Huang, Po-Yao Wang, Chieh-Chen Huang

**Affiliations:** Department of Life Sciences, National Chung Hsing University, Taichung 402, Taiwan; Innovation and Development Center of Sustainable Agriculture, National Chung Hsing University, Taichung 402, Taiwan; Advanced Plant and Food Crop Biotechnology Center, National Chung Hsing University, Taichung 402, Taiwan; Doctoral Program in Microbial Genomics, National Chung Hsing University and Academia Sinica, Taichung City and Taipei City 402 and 115, Taiwan; Department of Food Science and Biotechnology, National Chung Hsing University, Tai chung City 402, Taiwan

## Abstract

Several metabolites within the reductive tricarboxylic acid (rTCA) cycle have been found to form prebiotically. However, how these metabolites connect to each other and form rTCA cycle remains unresolved. The rTCA cycle is an ancient route and is considered significant for the emergence of life, since it connects to the routes of amino acids and nucleobases synthesis. A major challenge to complete the rTCA cycle under prebiotic conditions is the thermodynamically unfavorable reductive carboxylation of succinate to α-ketoglutarate. Here, we address this challenge by using the nature of energy: nonequilibrium conditions.

By calculating the changes in free energy, Δ*G*, of succinate to α-ketoglutarate, and its downstream reactions: α-ketoglutarate to glutamate and α-ketoglutarate to isocitrate under different nonequilibrium conditions, we find that these two-step reactions are exergonic under nonequilibrium conditions at a 10000:1 reactant-to-product ratio at 1.013 bar, pH 10 and 70°C. To prove the concept, we catalyze succinate to glutamate at a 10000:1 reactant-to-product ratio, with NH_2_OH and sodium dithionite. The process is catalyzed by Fe(0), Fe_3_O_4_, and artificial proto-[4Fe4S] clusters in 1M NaCl at pH 10 and 70°C under 1 atm of ^13^CO_2_ for 48 hours.

This nonequilibrium condition and one-pot system successfully promote the formation of α-ketoglutarate through carbon fixation with succinate and its subsequent conversion to glutamate. These findings demonstrate nonequilibrium states enable α-ketoglutarate formation through succinate and CO_2_, and suggest that a tendency toward natural thermodynamics may serve as a driving force for autocatalysis in the origin of life.

**Importance:** How life began remains open, metabolism provides a key framework for origins. We use a simple and robust energetic principle to show that non-equilibrium conditions can drive the highly endergonic carboxylation step of the reverse tricarboxylic acid (rTCA) cycle, enabling one-pot synthesis of glutamate.

This is work bridges the gap between protometabolites and protometabolsim, suggesting that metabolites may have accumulated first, creating concentration gradients that drove reactions and ultimately enabled the emergence of protometabolism. These findings provide a plausible pathway from prebiotic chemistry to the emergence of metabolism.

## Introduction

Life is a journey of electrons finding places to rest^1,2^. It manifests as complex, highly organized chemical networks to harness and release energy, involving the process of synthesis and breakdown^1,2^. The term protometabolism describes these self-sustaining networks as they exist in primitive environments^3^. This replenishable protometabolism was primarily driven by inorganic energy, including H_2_ and H_2_S. CO_2_ and small organic molecules present in the early Earth environment were materials building up fundamental building blocks in life^4–6^. Furthermore, minerals and metals in ancient geological environments functioned as catalysts, facilitating the synthesis of organic molecules from CO_2_^7–9^. Metabolism includes both catabolic and anabolic reactions, forming a nonequilibrium system that makes system continuously seek a new state of balance^3^. In extant research, 95% to 97% of the biochemical reactions were found exergonic under nonequilibrium state in mild alkaline environments when H_2_ served as a reductant^10^. Also, nonequilibrium conditions in cells are used to alter the ATP-to-ADP ratio, thereby lowering the free energy for ATP hydrolysis. This type of ratio control can also determine the reaction flux proceeds through glycolysis or gluconeogenesis^11^, which is similar to Le Chatelier’s principle. These findings suggest that a nonequilibrium state could be used to alter direction in cells and even overcome the thermodynamics barrier.

Among existing metabolisms, the reductive tricarboxylic acid (rTCA) cycle is considered to be an ancient and significant route before the emergence of life. In early earth, rTCA cycle served as a replenishable pathway for synthesizing numerous building blocks, including amino acids, carbohydrates, and nucleotides with the influx of CO_2_ and energy. This process includes both reductive and oxidative direction^12^. Also, in modern microbes, chemolithoautotrophy mainly uses rTCA cycle as central metabolomic pathways^13^. In Current prebiotic research, it shows that five universal building blocks—acetate, pyruvate, oxaloacetate, succinate, and α-ketoglutarate—as well as four other rTCA metabolites, can be synthesized from pyruvate and glyoxylate. This process is catalyzed by Fe^2+^ under mild conditions without the existence of CO_2_. Additionally, in the presence of hydroxylamine, these reactions can lead to one-pot amino acid synthesis^14^. Meanwhile, pyruvate, which is the entry of rTCA cycle, can be synthesized from CO_2_ via a Wood–Ljungdahl (WL)-analogous pathway catalyzed by native iron, and magnetite. The formation of acetate, formate, methane and methanol have also been observed^15–17^. Moreover, from the thermodynamic perspective, the free energy of formation of each metabolite in rTCA cycle from CO_2_ and H_2_ is exergonic^18^, suggesting that these molecules can form rapidly and spontaneously. However, these results are more like a one-way synthesis pathway which is different from metabolism. It only indicates that the formation of metabolites in prebiotic conditions is feasible^3^.

However, to run through the complete rTCA cycle under prebiotic environment is a challenge due to the high energy barrier. Among the rTCA cycle, there are five metabolites considered to be universal which facilitate three key carbon assimilation reactions: (1) acetate to pyruvate, (2) pyruvate to oxaloacetate, and (3) succinate to α-ketoglutarate. Among these, the reductive carboxylation reaction: succinate to α-ketoglutarate is the most endergonic and thermodynamically unfavorable reaction^18,19^. This reaction is also the gateway to glutamate synthesis, and then leads to glutamine, proline, and arginine synthesis^20^. In extant biology, α-ketoglutarate: ferredoxin oxidoreductase (KOR) catalyzes the carboxylation of succinyl-CoA and CO_2_ to form α-ketoglutarate. *KOR* genes are highly conserved among deep-branching microbial groups and are widely utilized for CO_2_ assimilation in autotrophic bacteria inhabiting extreme environments^21,22^. Interestingly, KOR in a deep-branching bacterium, *Aquifex Aeolicus*, is homologous with pyruvate: ferredoxin oxidoreductase (PFOR), a reaction catalyzes acetate to pyruvate. This suggests that in ancient microbes, they use one enzyme to catalyze two reductive carboxylation reactions: acetate to pyruvate and succinate to α-ketoglutarate^23^. Our previous study also demonstrated that the heterologous expression of *Chlorobium tepidum* KOR alone enables *Escherichia coli* to maintain cellular function under chemolithotrophic conditions^24^, highlighting the critical role of KOR as the primary CO_2_ entry point of the metabolic network for autotrophic growth.

While various prebiotic metabolites, including amino acids, have been experimentally shown to form under prebiotic conditions, it remains unclear how these metabolites within the rTCA cycle initially became interconnected.. The carboxylation of succinate to ***α***-ketoglutarate is particularly significant, as it represents the cycle’s most endergonic and thermodynamically unfavorable step. In modern cells, ATP, ferredoxin and coenzyme A are essential to catalyze the reaction^21^. In the reductive carboxylation step, [4Fe4S] clusters are commonly required and serve as the catalytic center in the carbon fixation protein^21,22^, and act as electronic capacitors to trap and release electrons in the carboxylation step^25^. Therefore, [4Fe4S] clusters have been considered a key element for the last universal common ancestor (LUCA)^26^, an essential role in the emergence of life^27^. In current prebiotic chemistry, it has been discovered that using sulfide, Fe^2+^, with a low concentration of cysteine in an anaerobic and alkaline environment the proto-[4Fe4S] cluster could form spontaneously^28^. Cysteine can also be prebiotically synthesized from serine in a mild environment^29,30^. Additionally, [4Fe4S] could also be synthesized by UV lights^31^.

In this study, we integrated a nonequilibrium system to cooperate the anabolism and catabolism into system, to see whether the most endergonic step of the rTCA cycle: succinate to α-ketoglutarate can thus go through. To do so, we coupled succinate to α-ketoglutarate with its downstream reaction to construct a nonequilibrium system. Magnetite (Fe_3_O_4_), native iron (Fe(0)), and an artificial proto-[4Fe4S] cluster were used as catalysts and electron capacitors. Success in driving this reductive carboxylation suggests that metabolites may have emerged first, establishing concentration gradients that facilitated further interconnections under the H_2_, CO_2_, and energy-rich conditions of early Earth. Such processes may have evolved into diverse metabolic pathways and, eventually, a self-replenishing network—a complete rTCA cycle that, like proto-metabolism, encompasses both anabolism and catabolism.

## Results

### The Gibbs Free Energy of Succinate to α-Ketoglutarate Reaction and Its Downstream Reaction under Nonequilibrium States

The eQuilibrator API was utilized to analyze the Gibbs free energy (ΔG) of reactions within the reductive tricarboxylic acid (rTCA) cycle under various nonequilibrium conditions. The analysis focused on the initial reductive carboxylation step, succinate to *α*-ketoglutarate, and two key downstream reactions: *α*-ketoglutarate to glutamate and *α*-ketoglutarate to isocitrate. Nonequilibrium states were simulated by adjusting the concentration ratios of the primary organic reactants and products. For example, succinate + CO_2_ + H_2_ ⇋α-ketoglutarate + H_2_O, the concentration ratio of succinate and α-ketoglutarate was adjusted to 100:1, 1:1, and 1:100 to generate different equilibrium states.

Based on the calculations, results show that the reductive carboxylation reaction of succinate to α-ketoglutarate remains endergonic, regardless of the reactant-to-product ratios being at 100:1, 1:1, or 1:100. However, the free energy dropped about 13kJ/mol and 27kJ/mol at 100:1 (reactant-to-product), compared with the reactant-to-product ratios being at 1:1 and 1: 100, respectively. As for the downstream reactions, α-ketoglutarate to glutamate and α-ketoglutarate to isocitrate, both reactions in three different nonequilibrium states were exergonic. Furthermore, the most exergonic value occurs at a 100:1 ratio, where the ΔG values were −53.5kJ/mol and −48kJ/mol in the reaction α-ketoglutarate to glutamate and α-ketoglutarate to isocitrate. As for the ratios at 1:1 and 1:100, the ΔG values were −40.3kJ/mol and −35kJ/mol for the α-ketoglutarate to glutamate reaction, and −27.2kJ/mol and −22kJ/mol for the α-ketoglutarate to isocitrate reaction (Fig. 1a). Though the succinate to α-ketoglutarate reaction persists as a non-spontaneous reaction, the influence of a nonequilibrium state on the Gibbs free energy of all reactions can still be observed.

**Fig 1.**
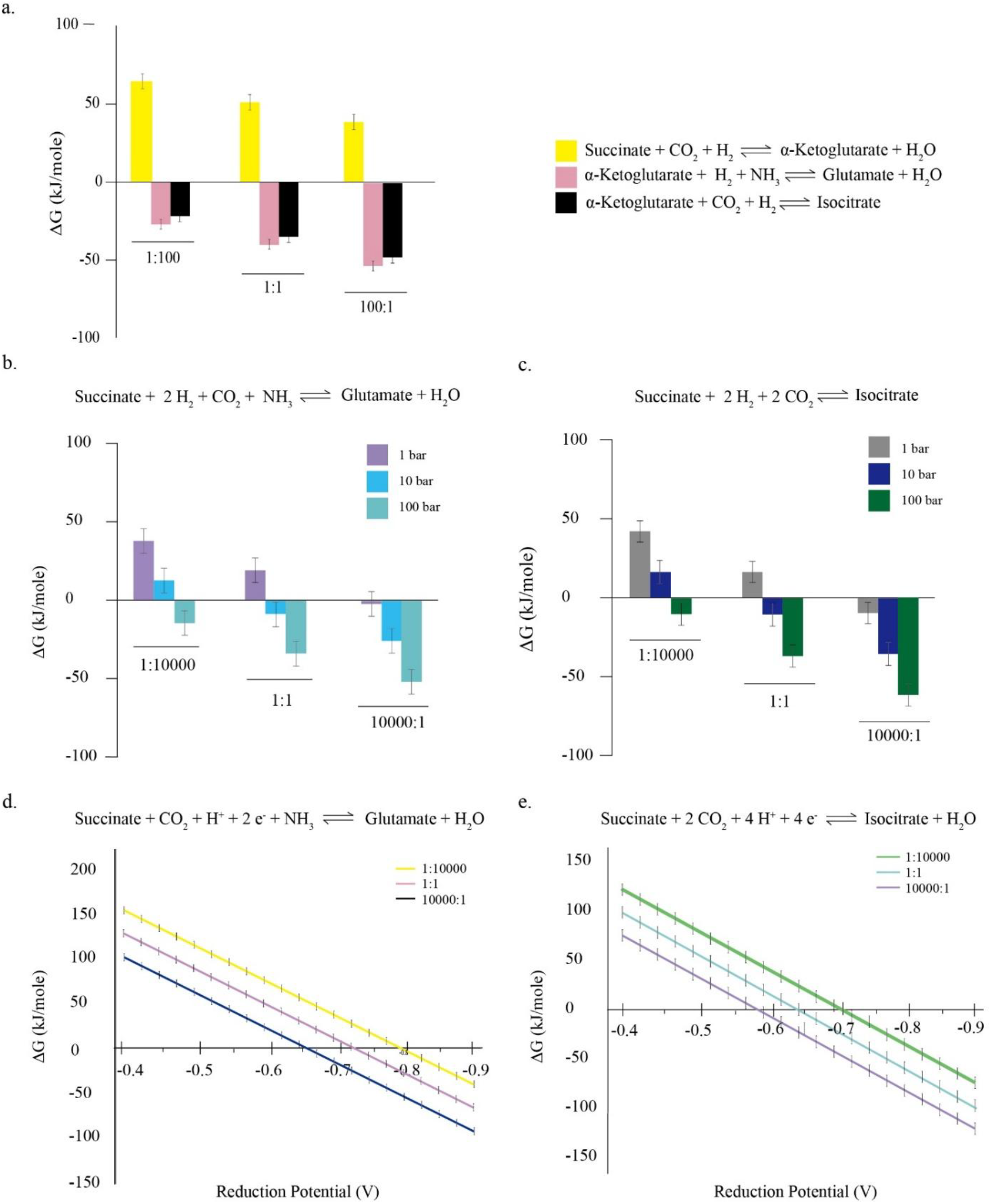
Thermodynamics calculation using the eQuilibrator API at different reactant and product ratios. In all reactions, 100:1 means the reactant and the product concentration were set to 10mM and 100μM, 1:1 means reactants and products were 10mM, and 1:100 indicates the reactant was 100μM and the product was 10mM. The concentrations of H_2_, CO_2_, and NH_3_, they were fixed at 0.64mM, 10mM, and 10mM, respectively. The conditions were set at 70°C (343.15K), pH=10, and ionic strength 1M. (a) The Gibbs Free energy of different reactions under different equilibrium states under 1bar. (b) The Gibbs Free energy of the succinate to glutamate reaction under different pressures and equilibrium states. (c) The Gibbs Free energy of the succinate to isocitrate reaction under different pressures and equilibrium states. (d) The reduction potential and the Gibbs free energy of the succinate to glutamate reaction at different concentration ratios. (e) The reduction potential and the Gibbs free energy of the succinate to isocitrate reaction at different concentration ratios.

### The Gibbs Free Energy of the Two-Step Combination Reaction Containing Succinate to α-Ketoglutarate Reaction and its Downstream Reaction of the rTCA Cycle under Nonequilibrium States

Based on the negative Gibbs free energy of the downstream reactions, we then expanded the system to two-step reactions by combining each downstream reaction with the succinate to α-ketoglutarate reaction. The reactions of succinate to glutamate and succinate to isocitrate were examined. To maintain a 100-fold difference among the concentrations of the reactant, intermediate, and product, the reactant concentration was set to 1M, and the product concentration was 100μM, having a highly unbalanced ratio of 10000:1. Conversely, under the 1:10000 conditions, the concentrations of reactant and product were reversed. For the 1:1 condition, both reactant and product were set to 10 mM.

The ΔG values of the two-step combination reactions were −15.5kJ/mol for the succinate to glutamate reaction and −10 kJ/mol for the succinate to isocitrate reaction at 10000:1 (Fig. 1b, 1c). These indicate that when the succinate to α-ketoglutarate reaction is combined with downstream reactions into two-step reactions, both two-step combination reactions are exergonic at a 10000:1 ratio under 70°C, pH=10, 1M ionic strength and, 1 bar conditions. The results show that the reaction toward downstream product synthesis and having a nonequilibrium state is thermodynamically favorable. As for the reaction ratios that are 1:1, and 1:10000, both two-step combination reactions, succinate to glutamate and succinate to isocitrate, remain endergonic at 1 bar atmosphere. Next, to see how pressure can influence the ΔG values under different nonequilibrium states, we increased the pressure of the system to get closer to the environment of the origin of life, where the alkaline hydrothermal vents were found at a water depth of 700m^5^. The pressurized system shows that the reaction becomes more exergonic in the 10000:1 ratio group, with the ΔG value even up to −50kJ/mol at 100 bar. Moreover, the reactions at a 1:1 reactant-to-product ratio was also exergonic at 10 and 100 bar. At pressures above 100 bar, the system at 1:10000 also becomes exergonic.

### The Gibbs Free Energy of the Two-Step Combination Reaction with Different Reduction Potentials under Nonequilibrium States

The reductive carboxylation step: succinate to α-ketoglutarate is the key route toward amino acid synthesis, which can lead to the synthesis of glutamate, glutamine, proline, and arginine, subsequently. Although the reaction is exergonic when H_2_ is served as an electron donor in the reaction at a 10000:1 ratio under 1 bar, the high energy barrier still poses a significant kinetic challenge to the reductive carboxylation step, the ΔG value is merely below zero. We then investigate the relationship between reduction potential and the Gibbs free energy.

The reactions: succinate + 4e^-^ + 4H^+^ + NH_3_ + CO_2_ ⇋Glutamate + H_2_O, and succinate + 4e^-^ + 4H^+^ + 2CO_2_ ⇋Isocitrate + H_2_O were calculated. From our results, when the potential energies from succinate to glutamate and succinate to isocitrate are below −650 mV and −587.5 mV, the reactions become exergonic at a reactant-to-product ratio of 10000:1 under 1bar. While decreasing the concentration of reactant, or increasing the concentration of product, the thresholds for proceeding reaction spontaneously become increasingly high (Fig. 1d, 1e).

In our system, the reduction potential of H_2_ is approximately −680 mV. The above result shows that the ΔG value of the reaction becomes slightly thermodynamically favorable at a 10000:1 reactant-to-product ratio. In modern bacteria, the KOR enzyme catalyzes succinyl-CoA to α-ketoglutarate. The reduction potential of Ferredoxin (Fd) in KOR is around −500 to −635 mV. This is achieved by using CoA, thiamine pyrophosphate, and an enzyme structure to decrease the energy barrier. The results so far show that the nonequilibrium state has a great impact on decreasing the reduction potential of the reaction and displays a thermodynamically favorable effect, which indicates the reductive carboxylation step, succinate to α-ketoglutarate, might occur together with downstream reactions under nonequilibrium states.

To test the modelling results, we used the succinate to glutamate reaction at a 10000:1 ratio as our target since the conversion of α-ketoglutarate to glutamate has been achieved using Fe(0) at 70°C with NH_2_OH in previous research^13^. In the following experiments, we applied two different types of reducers, H_2_ and sodium dithionite, one of which is the reducer commonly used in anaerobic bacteria and abundant in alkaline hydrothermal vents, and the other is a strong reducer that is usually used as an electron donor to [4Fe4S] clusters for the enzyme essay. In addition, we used artificial proto-[4Fe4S] clusters as electron capacitors for reductive carboxylation. As for the catalysts, Fe(0) and Fe_3_O_4_ were used in our system, given their ability to catalyze CO_2_ fixation into pyruvate and methanol and their abundance on early Earth^14^.

### Carbon Fixation Reaction under a Nonequilibrium State with Hydrogen as the Electron Donor

From the above thermodynamics modeling results, succinate to glutamate is feasible under a nonequilibrium state at 70ºC, pH=10, 1.013 bar, which include the synthesis of α-ketoglutarate within the two-step reaction. First, to ensure glutamate can also be synthesized from α-ketoglutarate our experiment, α-ketoglutarate and NH_2_OH were catalyzed by Fe(0), Fe_3_O_4_, and artificial proto-[4Fe4S] clusters at 70ºC, pH=10, 1.013 bar with 1M NaCl, and purged with ^13^CO_2_ and H_2_ (CO_2_: H_2_= 2:8), for 48hrs. The High-Performance Liquid Chromatography-Tandem Mass Spectrometry (HPLC-MS/MS) was used to analyze the outcome and compared with authentic standards (Supplementary Fig. S1). Glutamate (m/z 148) was detected through HPLC-MS/MS, indicating glutamate can be successfully synthesized from α-ketoglutarate under such environment (Supplementary Fig. S4). After making sure that glutamate synthesis can synthesized through reductive amination from α-ketoglutarate, we then investigated CO_2_ fixation from 1M succinate with 100μM glutamate following the ratio 10000:1, and catalyzed by Fe_3_O_4_, and Fe(0) with freshly synthesized artificial proto-[4Fe4S] clusters, NH_2_OH, and purged with ^13^CO_2_ and H_2_ (^13^CO_2_: H_2_= 2:8) at 70°C, pH=10, 1M NaCl for 48hr (Fig. 2a), comparing with the control groups that were purged with CO_2_ and H_2_.

**Figure 2.**
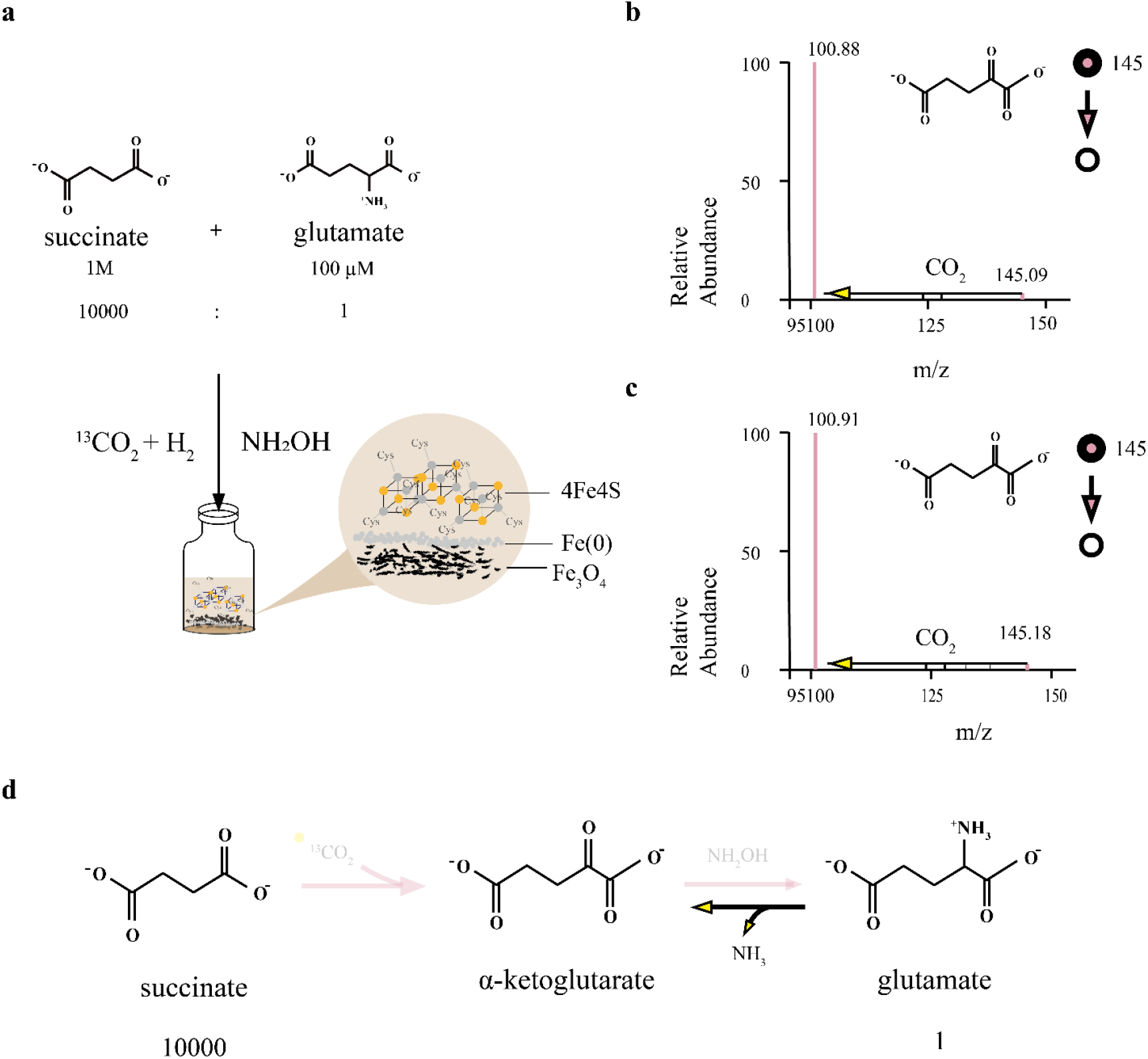
The Experiment design and results. (a) Experiment scheme. 1M succinate, 100µM glutamate, and 14mg hydroxylamine catalyzed by Fe(0), Fe_3_O_4_, and 4Fe4S purged with ^13^CO_2_ and H_2_ (CO_2_: H_2_= 2:8) at 70 ºC, pH=10, and 1 atm with 1M NaCl for 48hr. (b) (c) The LC-MS/MS results of (a) at 1.013 bar and 20 bar. (d) The hypothesized pathway.

After 48 hours, α-ketoglutarate was detected by HPLC-MS/MS, having ion mass m/z 145 to m/z 101, which corresponds to authentic standards (Supplementary Fig. S1), both shown in the experimental group and the control group (Fig. 2b, c, and Supplementary Figure S6). However, no M+1 α-ketoglutarate (if carboxylation happened from succinate, α-ketoglutarate would incorporate one ^13^C atom, m/z will be 146) has been detected through product ion scanning mode, regardless of 1 bar or 20 bar pressure in both the experimental group and the control group. These suggest the production of α-ketoglutarate might come from glutamate through deamination instead of the reductive carboxylation reaction from succinate with ^13^CO_2_ (Fig. 2d) and the concentration of α-ketoglutarate might be very scarce. Based on the results, it it possible that H_2_ might not be strong enough for catalyzing succinate to glutamate under a nonequilibrium environment in this experiment, albeit in 20 bars. Therefore, a stronger reducer might be needed. Later, we then substitute H_2_ with a strong reducer, sodium dithionite.

### Carbon Fixation under a Nonequilibrium State promoted by sodium dithionite as electron donor

We replaced H_2_ with sodium dithionite, which is a stronger reducing agent having a reduction potential of around −660 mV at standard conditions. Having made sure that glutamate can also synthesized through α-ketoglutarate when sodium dithionite as an electron donor under the same conditions (Supplementary Fig. S5), we then use sodium dithionite as a reducer in the following experiments: the two-step reaction catalysis involving reductive carboxylation reaction.

After 48 hours of catalysis, α-ketoglutarate was detected in the experimental groups purged with ^13^CO_2_. Both M+0 (m/z 145) and M+1 (m/z 146) signals were detected. Among the m/z 146 signals, the m/z 146 → 101 fragment (Fig. 3a, Supplementary Figs. S7~8) was predominant to 106 → 102 fragment compared with the standard (Supplementary Figs. S1b, S2) and control groups which were purged with non-labeled CO_2_ (Supplementary Figs. S9). This indicates that most of the m/z 146 signal fragmented into m/z 45 (^13^CO_2_) and m/z 101, rather than into m/z 44 and m/z 102. Comparing with control groups which purged with non-labeled CO_2_, the m/z 146 → 101 fragments shown in experimental group demonstrating that the purging gas ^13^CO_2_ was incorporated into α-ketoglutarate. Additionally, non-labeled α-ketoglutarate (m/z 145) was also detected in the groups purged with ^13^CO_2_ (Supplementary Fig. S7). To analyze how non-labeled α-ketoglutarate were formed, we then check other potential inorganic carbon sources in our reaction. In the process of synthesizing artificial proto-[4Fe4S] clusters, NaHCO_3_ will be added to stabilize the structure. To evaluate whether NaHCO_3_ contributes to carbon incorporation, a control experiment was performed in which NaH^13^CO_3_ was used during cluster synthesis while the reaction system was purged with non-labeled CO_2_. HPLC–MS/MS analysis showed results identical to those of the control group which purged with non-labeled CO_2_, indicating that NaHCO_3_ does not involve in carbon fixation reaction (Supplementary Fig. S10). This indicates that the non-labeled α-ketoglutarate (m/z 145) shown in the experimental groups which purged with ^13^CO_2_ might come from deamination reaction from glutamate. These results suggest that both forward and reverse pathways occurred within 48 h, including reductive carboxylation to α-ketoglutarate and deamination of glutamate to α-ketoglutarate.

**Figure 3.**
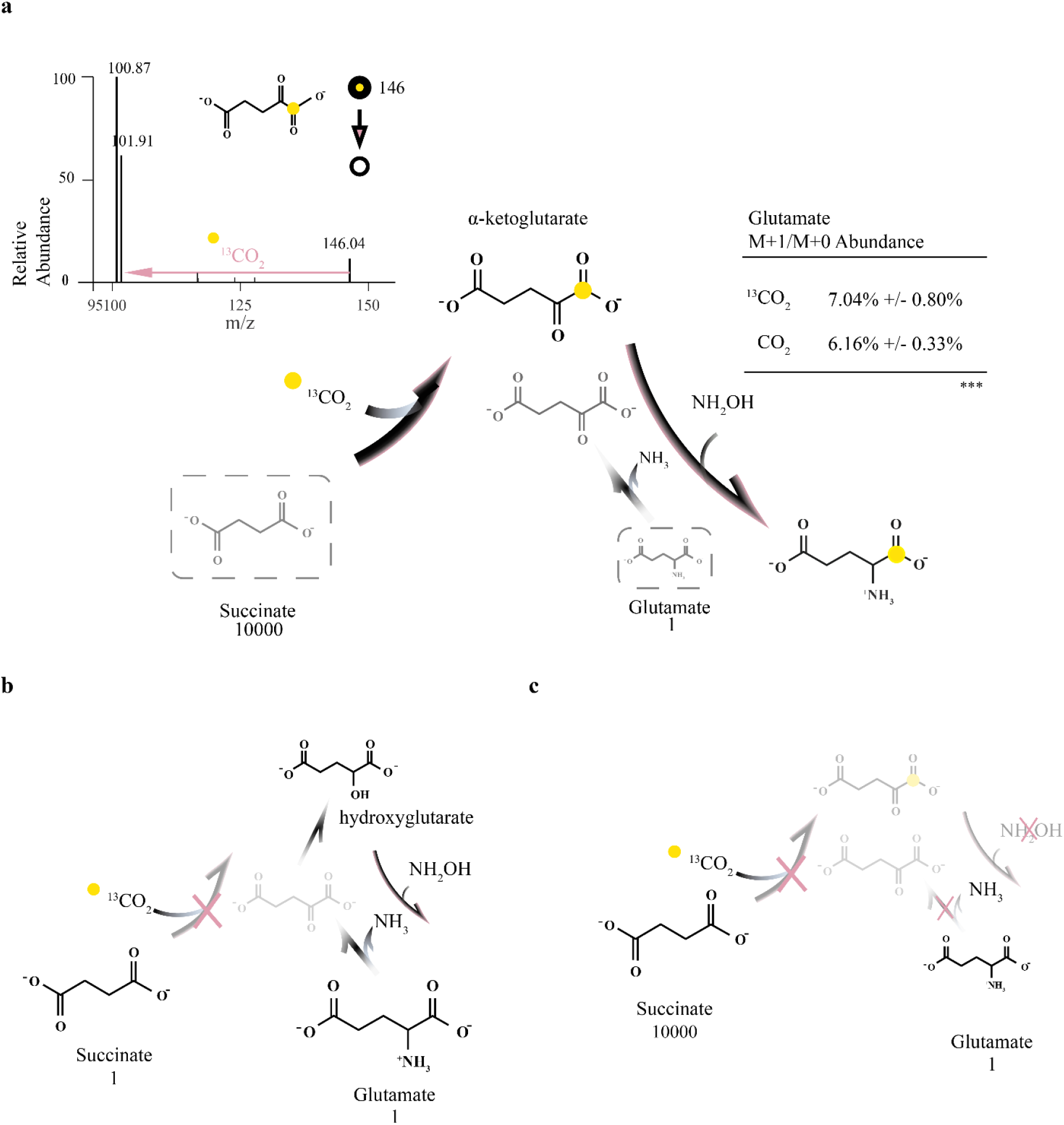
Nonequilibrium state promotes carbon fixation reaction through two step catalysis. (a) The HPLC-MS/MS result of 1M succinate, 100µM glutamate, 14mg hydroxylamine, and 34.8mg sodium dithionite, catalyzed by Fe(0), Fe_3_O_4_, and 4Fe4S, purged with ^13^CO_2_ at 70 ºC, pH=10, and 1 atm with 1M NaCl for 48 hours. ***means p < 0.05. (b) The result of reactant succinate and glutamate are under same concentration ratio (1:1). (c) The result of reaction without adding hydroxylamine.

Further, to know whether the α-ketoglutarate formed from succinate with ^13^CO_2_ could be synthesized into glutamate, the M+1/M+0 ratios of glutamate in the same samples from the experimental groups and the control groups were analyzed. 7.04% of M+1/ M+0 glutamate ratio was shown in experimental groups purged with ^13^CO_2_ compared with the 6.16% of M+1/M+0 glutamate ratio in the control group purged with CO_2_ (Fig. 3a, Supplementary Table 1, 2). Despite the modest ~1% increase, the difference is statistically significant. These two HPLC-MS/MS results from α-ketoglutarate and glutamate indicate that the two step conversion of succinate to glutamate can occur under nonequilibrium condition.

In addition to α-ketoglutarate, isocitrate, glyoxylate, pyruvate, and malate were also examined in experimental groups. However, no corresponding peaks were detected for these compounds. Notably, prominent peaks at m/z 191 and m/z 193 were detected in both the experimental groups purged with ^13^CO_2_ and the control group purged with CO_2_ (Supplementary Fig. S13), however the peaks pattern differed from those of the standard isocitrate compound (Supplementary Fig. S1d). This suggests that these peaks are unlikely to be isocitrate.

To make sure that the nonequilibrium condition is the key factor for driving carbon fixation to happen, we investigated the reactions under reactant-to-product (succinate: glutamate) concentration ratios at 1:1. After 48 h, neither non-labeled α-ketoglutarate nor labeled α-ketoglutarate was detected. In contrast, hydroxyglutarate was detected by LC–MS/MS (Fig 3b, Supplementary Fig. S11). Furthermore, reactions without adding NH_2_OH were also analyzed, non-labeled α-ketoglutarate, labeled α-ketoglutarate, and hydroxyglutarate were not detected by HPLC-MS/MS (Fig. 3c, Supplementary Table 3), which results were as same as the group without adding metals, and artificial proto-[4Fe4S] clusters. These results suggest the necessities of NH_2_OH and minerals.

Though the nonequilibrium reaction cannot go further to isocitrate or other metabolites in the rTCA cycle, these results still represent that carbon fixation under nonequilibrium states α-ketoglutarate can achieved through reductive carboxylation from succinate and deamination from glutamate with NH_2_OH, sodium dithionite, catalyzed by Fe_3_O_4_, Fe(0)and artificial proto-[4Fe4S] clusters. Remarkably, in the results, labelled-α-ketoglutarate can also lead to glutamate synthesis in 48 hours.

### Carbon Fixation Reaction under Nonequilibrium States with different metal combinations

To know the necessity of metals and artificial proto-[4Fe4S] clusters, different combinations were analyzed for their abilities to drive carbon fixation. The reactions were carried out in the same condition with sodium dithionite, NH_2_OH, 1M succinate, and 100μM glutamate purged for CO_2_ or ^13^CO_2_ at 70°C, 1M NaCl, pH 10, and reaction time for 48 hours under 1.013 bar. 145 m/z to 101 m/z and 146 m/z to 101 m/z were analyzed by HPLC-MS/MS in all groups to see whether α-ketoglutarate and labeled α-ketoglutarate were produced. In our result, 146 m/z (M+1) to 101 m/z α-ketoglutarate was detected only in groups 2 and 3 (Fig. 4a, Supplementary Fig. S12), albeit M+1 α-ketoglutarate was detected in one of three replicates in group 3.

**Figure 4.**
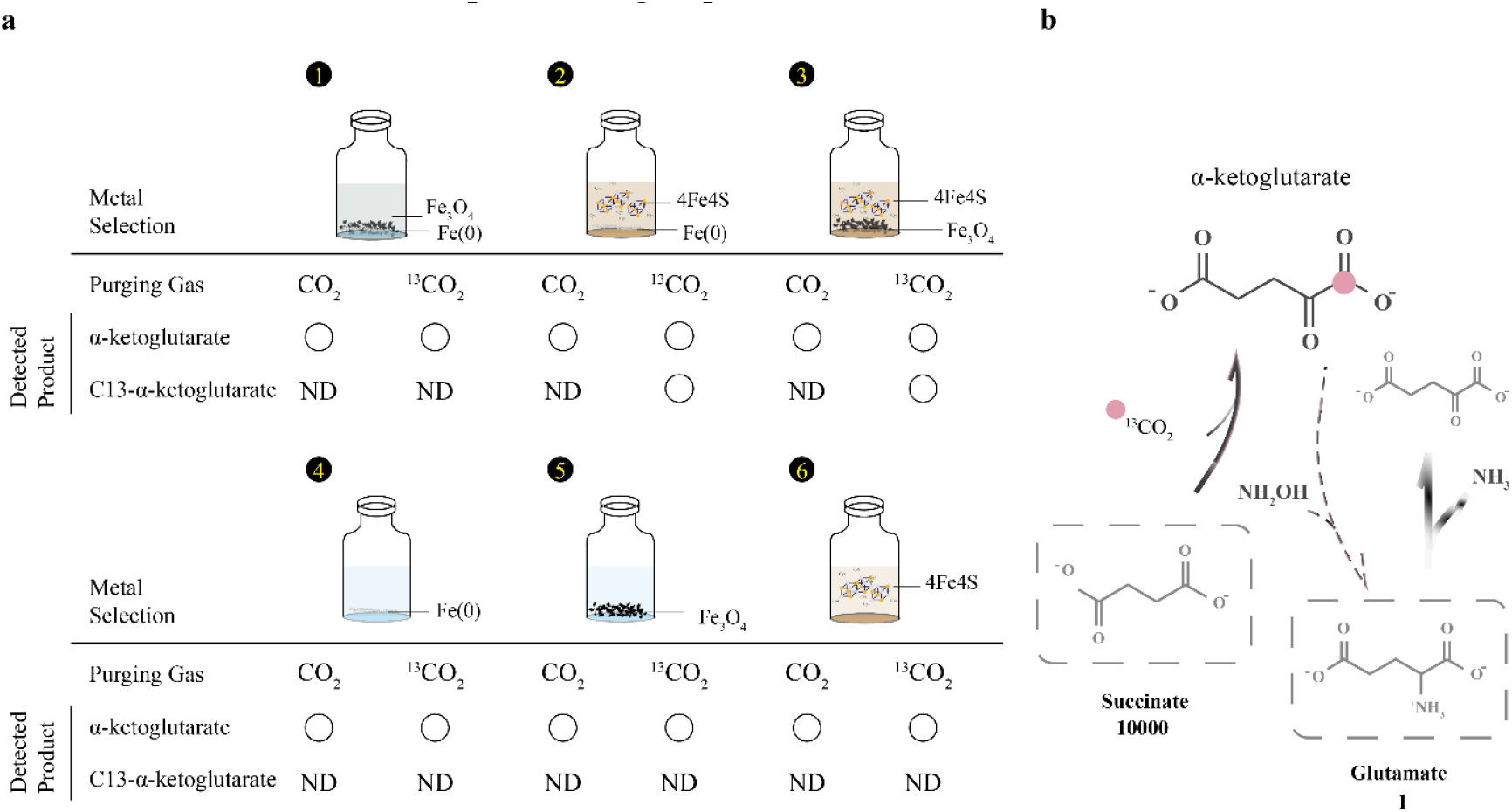
Carbon Fixation Reaction under nonequilibrium states with metal combinations. (a) The HPLC-MS/MS result of the reaction catalyzed by different metal combinations. (b) the result scheme of the reaction catalyzed by 4Fe4S with Fe_3_O_4_ and 4Fe4S with Fe(0).

By comparing groups 4 and 5 with groups 2 and 3, it is evident that the artificial proto-[4Fe4S] cluster is essential in the carbon fixation reactions. Similarly, comparison between group 1 with the experiment previously catalyzed by Fe_3_O_4_, Fe(0), and artificial proto-[4Fe4S] clusters (Fig 3), also emphasize the significant catalytic role of artificial proto-[4Fe4S] clusters. In the group 6, the presence of artificial proto-[4Fe–4S] clusters alone was insufficient to successfully catalyze reductive carbon fixation, suggesting the importance of Fe(0) and Fe_3_O_4_.

## Discussion

In this research, we observed the two-step reaction: succinate to glutamate can be achieved under nonequilibrium condition while sodium dithionite as electron donor, reactant-to-product ratio at 10000:1 with Fe(0), Fe_3_O_4_ and artificial proto-[4Fe4S] clusters as catalysts at 1m NaCl, pH=10, 70°C, 1atm, and purged with ^13^CO_2_, reaction time for 48hr. When comparing the effects of different electron donor, sodium dithionite can drive carbon fixation reaction toward glutamate synthesis, with Fe(0), Fe_3_O_4_, and artificial proto-[4Fe4S] clusters at 70°C, pH 10, 1M NaCl under nonequilibrium states (10000:1), whereas no α-ketoglutarate signal synthesized from succinate and ^13^CO_2_ was detected, regardless of 1.013 or 20 bar pressure when H_2_ served as reductant.

Hydrogen was widely considered to be an energy source in the primitive environment. In prebiotic chemistry, H_2_ and CO_2_ can be converted into formate, acetate, and pyruvate in the presence of Fe(0) or Fe_3_O_4_ through a highly exergonic pathway^4,13,14^. Also, in modern biology, many autotrophs derive their energy from H_2_. A study illustrated that H_2_ cannot catalyze ferredoxin without the presence of Fe(0)^32^. The midpoint potential approximately −414 mV of H_2_ under physiological conditions is insufficient to reduce the ferredoxin, whose reduction potential is at −412mV. In our system, artificial proto-[4Fe4S] clusters are considered essential in the reductive carboxylation reaction. However, the midpoint potential of artificial proto-[4Fe4S] clusters at 70°C and pH 10 is unclear. The midpoint potential is around −400mV at room temperature with pH=10 according to previous research^28^. Whether H_2_ could reduce artificial proto-[4Fe4S] clusters still needs more experiments and could thus understand more about the role of artificial proto-[4Fe4S] clusters in the reaction. Also, from our thermodynamic calculations under nonequilibrium conditions (reactant-to-product ratio of 10000:1), the succinate to glutamate reaction requires a reduction potential lower than −650 mV at 70 °C and pH 10. The midpoint potential of H_2_ under the same conditions is approximately −670 mV^16^. This could indicate that H_2_ might have the ability to catalyze carbon fixation in a certain environment. However, the yield of α-ketoglutarate might be too scarce to be detected, or simply because the reduction potential of H_2_ is not enough for reducing artificial proto-[4Fe4S] clusters or catalyzing reductive carboxylation steps. Additionally, the low solubility of H_2_ should be taken into consideration.

To determine whether the two-step conversion from succinate to glutamate occurred due to nonequilibrium conditions or the high reducing power of sodium dithionite, we tested the reaction under reactant-to-product (succinate: glutamate) concentration at the same ratio. No M+1 α-ketoglutarate was detected. This reaffirming that it is the nonequilibrium state enabled the carbon fixation reaction, rather than the strong reducing potential of sodium dithionite. In the experiments, Fe(0), Fe_3_O_4_, and artificial proto-[4Fe4S] clusters were used as catalysts and electron capacitors. In previous research, the surface of Fe(0) and Fe_3_O_4_ could lower the activation energy for carbon fixation reactions. Moreover, Fe(0) can reduce water into H_2_^4,13^, providing H_2_ for the reaction in this research. As for [4Fe4S] cluster, it is used as electron carriers and is highly conserved in carbon fixation reductase^22,25^. In our metal selection experiments, for M+1 α-ketoglutarate production, it requires [4Fe4S] to couple with Fe(0) or Fe_3_O_4_, suggesting [4Fe4S] might play crucial role in this reaction, which might transfer the electron from sodium dithionite to the reaction, while Fe(0) or Fe_3_O_4_ served as the surface for reaction happened and reduced the activation energy of reaction.

In our modelling experiments, when pressure reaches 100 bar, the Gibbs free energy of succinate to glutamate is ~−50 kJ/mol. In the Lost City Hydrothermal Vent Field (LCHF), where the water depth is ~750 meters, the pressure is approximately 70 bar. The two-step reaction may be highly favorable at LCHF, particularly in an environment with abundant H_2_. Besides, ion gradients that are harnessed by chemiosmotic coupling may form within the pores or at the interface between the exterior and interior of the chimney structure. These ion gradients are considered an ancient mechanism analogous to ATP utilization in modern cells, where ATP serves as the primary energy currency and involves in reductive carboxylation reaction^9^. The ion gradients in LCHF might be the energy sources for two-step reaction. This suggests that reductive carboxylation reactions in primitive environments were highly favorable under nonequilibrium conditions.

In this research, we minimize potential background effects, all reactions were performed in triplicate under both CO_2_ and ^13^CO_2_ purging conditions. Also, we use product ion scan to double check the m/z 146 → 101 fragments and m/z 146 → 102 fragments, to ensure the m/z 146 → 101 in the experimental group is confidential (Supplementary Fig. S7, S8), compared to the control group which was purged with non-labeled CO_2_, and the control group which was added NaH^13^CO_3_. In this study, the M+1/M+0 ratio of glutamate was calculated and compared with the CO_2_-purged control group to determine whether ^13^C incorporation into glutamate had occurred. The M+1/M+0 ratio of glutamate was compared with its natural abundance in the standard and control groups (Supplementary Table 1, 2).

Here, we implement a relatively natural system, the nonequilibrium state, into the carbon fixation reaction to see whether succinate toward the synthesis of glutamate can drive a carbon fixation reaction (succinate to α-ketoglutarate). The result shows α-ketoglutarate successfully synthesized through carbon fixation with succinate and CO_2_ while sodium dithionite acted as reducer, catalyzed by Fe_3_O_4_, Fe(0) and artificial proto-[4Fe4S] clusters under a 10000:1 nonequilibrium system. Furthermore, in this 48hr, α-ketoglutarate synthesized from succinate and CO_2_ can further lead to glutamate synthesis in one-pot. The production of α-ketoglutarate, succinate, glutamate and other metabolites in rTCA cycle have been shown to be easily and rapidly formed by adding glyoxylate with pyruvate and catalyzed by Fe^2+^ at 70°C with amino acids can be synthesized in one-pot by adding NH_2_OH^3^. Moreover, pyruvate can be formed through CO_2_ catalyzed by Fe(0) or Fe_3_O_4_^4^. With previous prebiotic research exhibits the feasibility of production of metabolites at primitive like environment. The result in our research might reveal that the rTCA cycle shows up after all the metabolites of the rTCA cycle and amino acids were formed. In this experiment, Fe(0) and Fe_3_O_4_, together with artificial proto-[4Fe–4S] clusters, enabled α-ketoglutarate synthesis under nonequilibrium conditions, supporting its prebiotic feasibility. These findings suggest that rTCA cycle metabolites could begin to interlink and form a replenishable network, once metabolites and amino acids are produced and a nonequilibrium state is established.

### Conclusion

We use a simple, natural, and robust concept to prove that the reductive carbon fixation step: succinate to α-ketoglutarate can be driven nonenzymatically under nonequilibrium states in the presence of low concentrations of glutamate in an alkaline-hydrothermal vent-like environment with Fe(0), Fe_3_O_4,_ and artificial proto-[4Fe4S] clusters, and also lead to glutamate synthesis in one-pot.

## Material and Methods

### Chemical

Succinate >=99%, glutamate, α-ketoglutarate >=98%, L-cysteine, FeCl_2_, Fe_3_O_4_ 97%, Na_2_S, and NaHCO_3_ >=99.7% were obtained from Sigma Aldrich (USA). Isocitrate 95% and Fe(0) 99% were purchased from Acros Organics. Sodium dithionite technical grade was purchased from sigma. Hydroxylamine hydrochloride 99% was purchased from Alfa Aesar. NaOH was obtained from Uni Region Biotech. NaCl <=100% was obtained from Honeywell Fluka. Carbon dioxide (^13^CO_2_) was purchased from Sigma Aldrich.

### Synthesis of proto-[4Fe4S] clusters

The synthesis procedure followed the method described by Jordan et al. (2021). NaHCO_3_ was added to the Na_2_S solution before adding L-cysteine and adjusting the pH to 10 with 1M NaOH. FeCl_2_·4H_2_O was then added into the solution, and the solution reached a final concentration of 10mM NaHCO_3_, 5mM cysteine, 1mM Na_2_S·9H_2_O, and 1mM FeCl_2_·4H_2_O with deaerated mili-Q water. All synthesis processes were carried out in an anaerobic chamber.

### Detection of proto-[4Fe4S] clusters

The freshly prepared solutions were analyzed through JASCO V-630 spectrophotometer for detecting 420nm UV-Vis absorption. The sample was placed into a plastic semi-micro cuvette inside the anaerobic chamber. The cuvette was then sealed with the lid before detection. All samples were recorded in triplicate immediately after preparation. All reactions for promoting carbon fixation used freshly prepared [4Fe4S] cluster, after the[4Fe4S] cluster having analyzed at 420nm.

### H_2_ as an Electron Donor for Promoting Reductive Carboxylation Reactions

All glassware was sunk in an acid bath overnight and cleaned with aqua regia if the vials were used with metals before. 2 mmol succinate, 0.0002 mmol glutamate, 2 mmol Fe(0), 2 mmol Fe_3_O_4_, 1/1000 of the artificial proto-[4Fe4S] cluster, and 2 mmol NaCl were placed into 10 ml glass vial. After pH was adjusted with 1M NaOH, glass vials were filled with deaerated milli-Q water to a final volume of 2 ml. 14 mg hydroxylamine hydrochloride powder were then added to the reaction vessels before the vessels were closed by caps with punctured PTFE septa and purged with H_2_ for 2 min. After purging, a syringe was used to withdraw H_2_ and inject ^13^CO_2_ to reach the expected volume ratio. For a pressurized reaction, four syringes were inserted into the PTFE septa of vials, and the vials were placed in a stainless steel 300 ml vessel immediately to reduce the contact with O_2_. It was then sealed and washed with N_2_ three times, then the steel was flushed with 4 bar ^13^CO_2_ and 16 bar H_2_. The steel was then heated to 70 ºC and 1000 rpm at incubator. All the reactions were operated in triplicate.

### Sodium Dithionite as an Electron Donor for Promoting Reductive Carboxylation Reactions

All glassware was sunk in an acid bath overnight and cleaned with aqua regia if the vials were used with metals before. 2 mmol succinate, 0.0002 mmol glutamate, 2 mmol Fe(0), 2 mmol Fe_3_O_4_, 1/1000 of the artificial proto-[4Fe4S] cluster, and 2 mmol NaCl were placed into 10 ml glass vial. After pH was adjusted with 1M NaOH, glass vials were filled with deaerated milli-Q water to a final volume of 2 ml. 35 mg sodium dithionite powder and 14 mg hydroxylamine hydrochloride powder were then added to the reaction vessels before the vessels were closed by caps with punctured PTFE septa and purged with N_2_ or ^13^CO_2_ for 2 min. The reaction vials were then placed into an incubator at 70ºC and 1000 rpm. All reactions were performed in an anaerobic chamber and were operated in triplicate.

### Reductive Amination Reaction from α-ketoglutarate Reactions

All glassware was sunk in an acid bath overnight and cleaned with aqua regia if the vials were used with metals before. 0.02 mmol α-ketoglutarate, 2 mmol Fe_3_O_4_, 1/1000 of the artificial proto-[4Fe4S] clusters, and 2 mmol NaCl were placed into 10 ml glass vial. After pH was adjusted with 1M NaOH, glass vials were filled with deaerated mili-Q water to a final volume of 2 ml. 35 mg sodium dithionite powders or purged with H_2_ for 2min, and 14 mg hydroxylamine hydrochloride powders were then added to reaction vessels before vessels were closed by caps with punctured PTFE septa and purged with N_2_ (if sodium dithionite was used as electron donor) for 2 min. The reaction vials were then placed into an incubator at 70ºC and 1000 rpm. All reactions were performed in an anaerobic chamber and were operated in triplicate.

### Sample Work-up Procedure

60 mg KOH was added into reaction vials and transferred into eppendorf in an anaerobic chamber. Centrifuge at 13,000 r.p.m for 10 min to precipitate the metals. (If no metals were added previously, then there’s no need to add KOH.) Then the supernatant was transferred to the 2 ml HPLC sample vials in an anaerobic chamber and capped with a PTFE septum. Stored the sample at −20 ◦ C before HPLC-MS/MS analysis.

### Analytical Methods for Reductive Carboxylation Reactions

An Agilent 1200 Series HPLC equipped with a HILIC column (TSKgel Amide-80 3 µm 150 mm x 2.0 mm; Tosoh Bioscience) and a LTQ XL™ Linear Ion Trap Mass Spectrometer (Thermo Fisher Scientific, United States) were used. Pure water with 0.05% NH_4_OH as solvent A, and acetonitrile with 0.05% NH_4_OH as solvent B were used. The gradient elution program was set as follows: Phase B was initially held at 70% for 1.5 min, then rapidly decreased to 55% over 0.1 min. It was maintained at 55% for 4.4 min, then increased back to 70% over 0.1 min, and held at 70% until 10 min. The flow velocity was set at 0.3 mL per minute. The negative ion mode parameter was set at 3.2kV spay voltage, 275 °C capillary temperature, 5 arb sheath gas flow rate, −27 V capillary voltage and −56.52 V tube lens. The positive ion mode parameter was set at 4.0kV spay voltage, 275 °C capillary temperature, 3 arb sheath gas flow rate, 2 arb sweep gas flow rate, 4 V capillary voltage and 35 V tube lens.

### Thermodynamic Calculation

Gibbs energies were calculated by eQuilibrator API 0.4.7 with Python 3.9.13. to calculate the change of Gibbs free energies, eQuilibrator uses the following equation to calculate the change of Gibbs free energies.

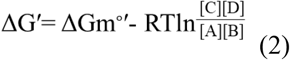

ΔGm^∘^′ means under a physiological environment pH 7, 25ºC, where reactants and products are at 1mM, and the reaction is at 1 atm gas pressure. R is the gas constant, T is the Temperature (K), [A], [B], [C], and [D] are the molar concentration of the reactant and product. ΔG′ is the change of Gibbs free energy under different temperatures and concentrations of reactant and product.

When at equilibrium condition, ΔG′=0, which means it will become equation (3), where the ratio of [A], [B], [C], and [D] will become equilibrium constant. In this study, the concentration of hydrogen was calculated by equation (4) based on the reference of H_2_ partial pressures and H_2_ concentrations in water at different temperatures^10^.

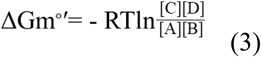

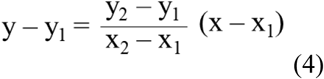

## Acknowledgments

This work was funded by National Science and Technology Council 113-2221-E-005-020-MY3 (Chieh-Chen Huang). We thank Dr. Chien-Chen Lai’s Lab for analyzing samples through HPLC-MS/MS, especially for Zi-Ji Lee. We thank Dr. Wen-Hua Chiou for providing the pressurized equipment. We thank Jui Teng Cheng from Dr. Wen-Hua Chiou’s lab for helping perform pressurized system. We thank Dong-Yan Wu and Shou-Chen Lo from Dr. Chieh-Chan Huang’s lab for technical suggestions. We thank Dr. En-Pei Isabel Chiang for helping with analysis in the beginning of the project. Finally, we especially thank Dr. Martina Preiner for reviewing the paper and giving meaningful suggestions.

## Author contributions

Conceptualization: Yu-Hsi Lin and Chieh-Chan Huang. Methodology: Yu-Hsi Lin, Jian-Hau Peng, and Shao-Yu Huang. Investigation: Yu-Hsi Lin, Po-Yao Wang. Validation: Yu-Hsi Lin. Formal analysis: Yu-Hsi Lin. Writing Original Draft: Yu-Hsi Lin. Review: Yu-Hsi Lin, Chieh-Chen Huang, Jian-Hau Peng, and Shao-Yu Huang. Visualization: Yu-Hsi Lin. Supervision: Chieh-Chen Huang. All authors reviewed and approved the final manuscript.

