## Supplementary Figure for "Nonequilibrium States Promote One-Pot Nonenzymatic Carbon Fixation in the Reverse Tricarboxylic Acid Cycle and Amino Acid Synthesis"

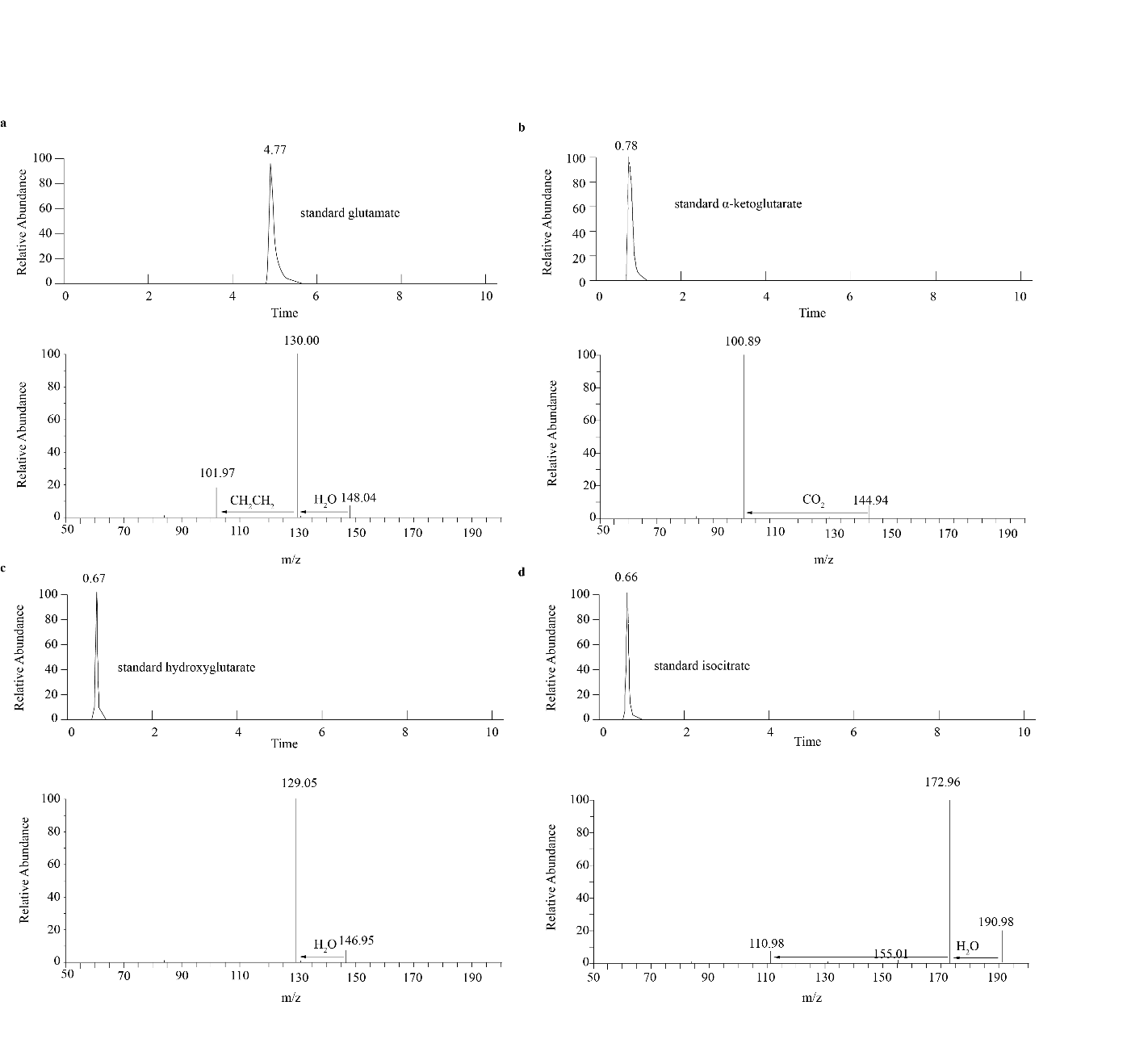

Figure S1. The HPLC spectra and MS/MS results of authentic standard. (a) the HPLC-MS/MS results of glutamate standard. (b) the HPLC-MS/MS results of α-ketoglutarate standard. (c) the HPLC-MS/MS results of hydroxyglutarate standard. (c) the HPLC-MS/MS results of isocitrate standard.

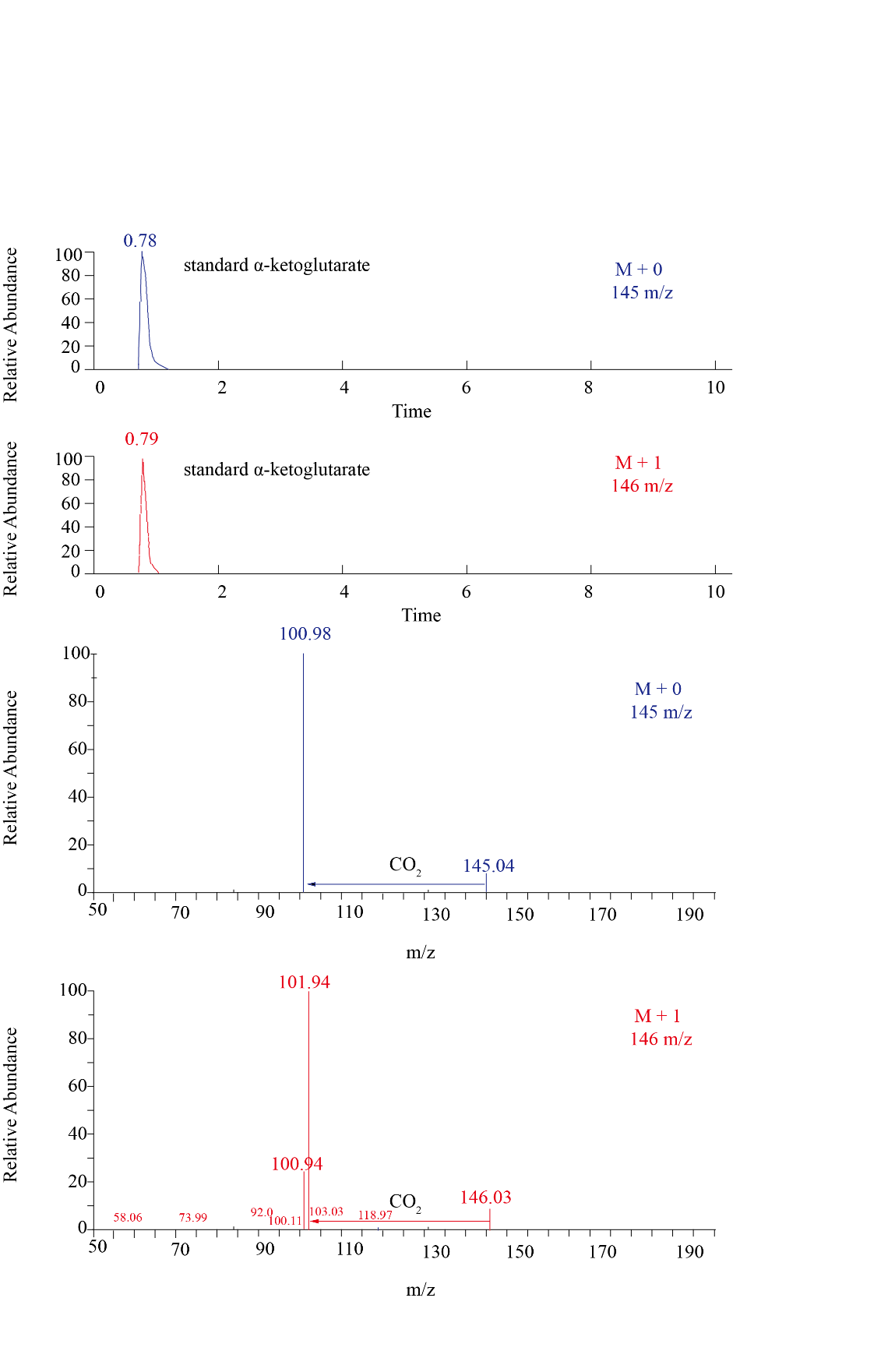

Figure S2. The HPLC spectra and MS/MS results of authentic α-ketoglutarate standard. Ion product scan at 100.5~101.5 m/z and 101.5~102.5 m/z from precursor ion m/z 145 (blue), and 146 (red), respectively.

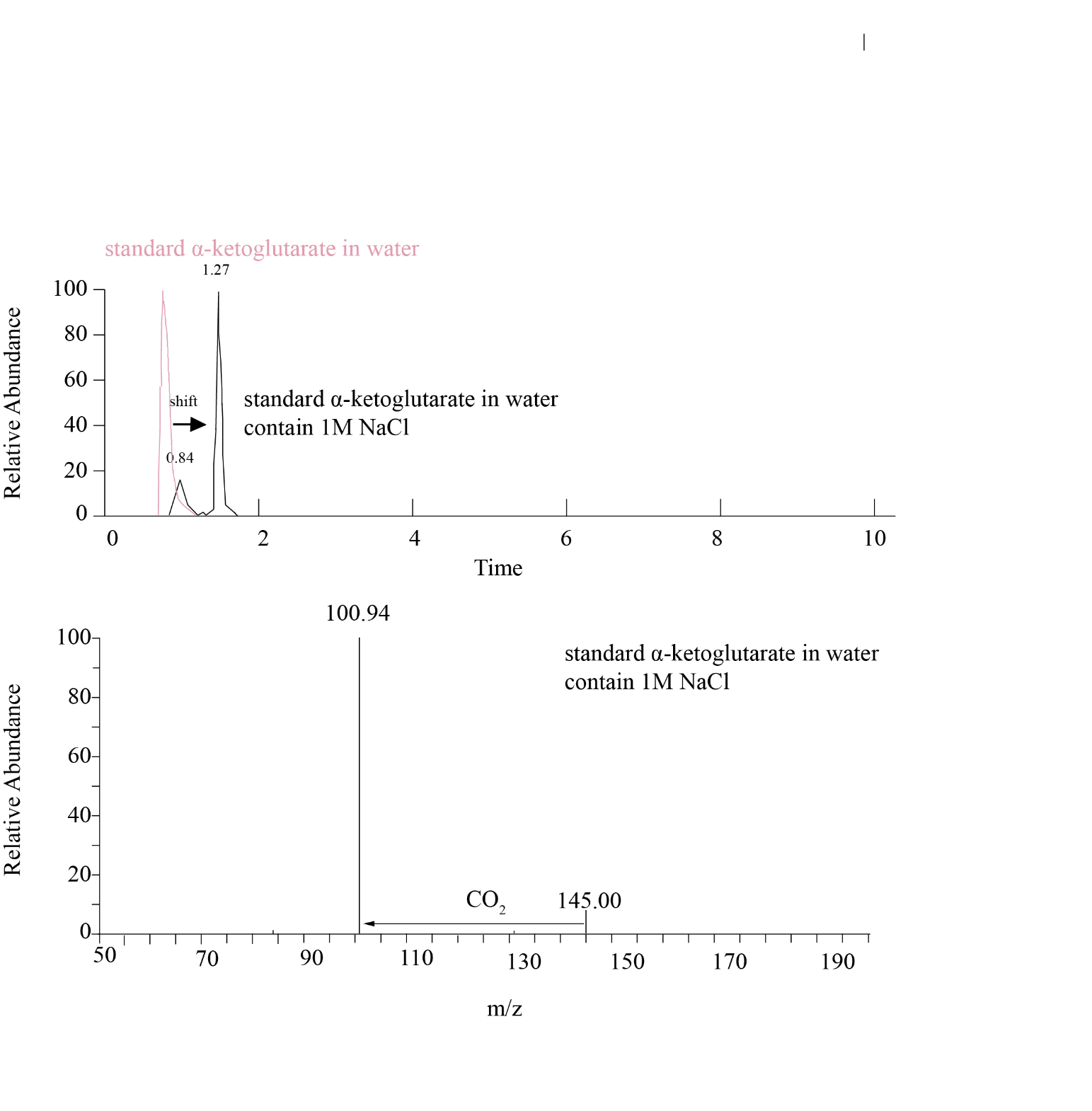

Figure S3. The HPLC spectra and MS/MS results of authentic standard -ketoglutarate in water pH=10 (pink line) and 1M NaCl water solution pH=10(black line).

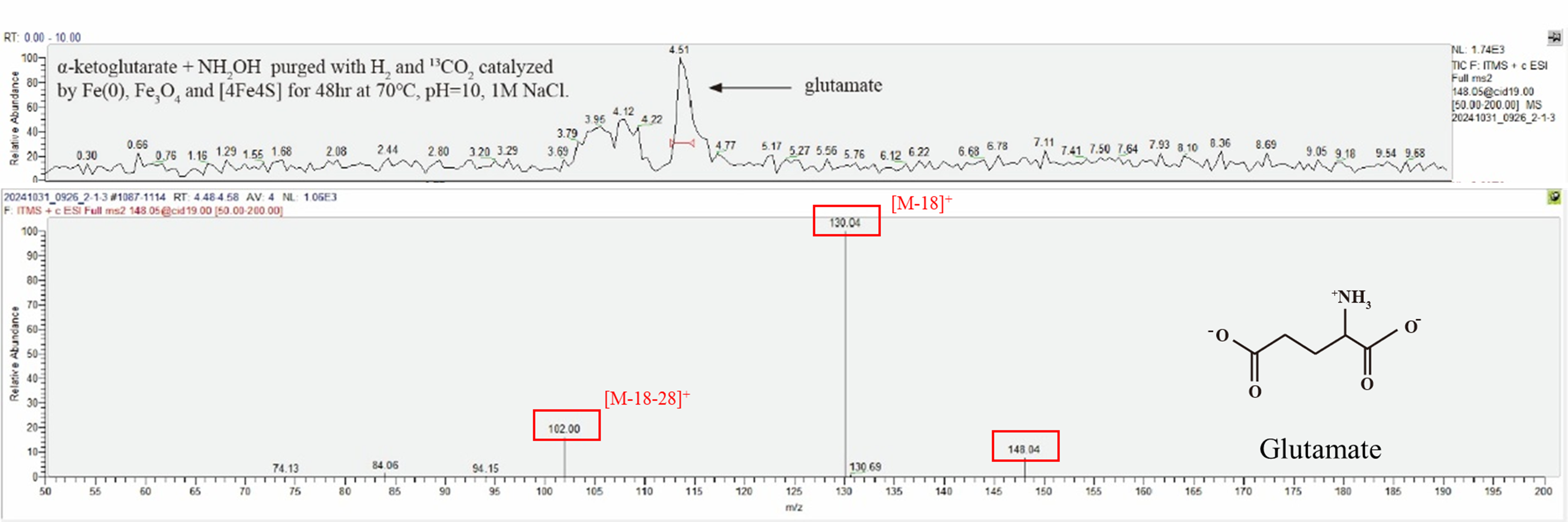

Figure S4. Glutamate synthesis from α-ketoglutarate, H_2_ as electron donor. HPLC–MS/MS spectra of m/z 148, the product reaction mixtures containing α-ketoglutarate and hydroxylamine, catalyzed by Fe(0), Fe₃O₄, and artificial [4Fe–4S] clusters with 1M NaCl. The reactions were purged with ^13^CO₂ and H₂ (2:8), then incubated at 70 °C for 48 hr.

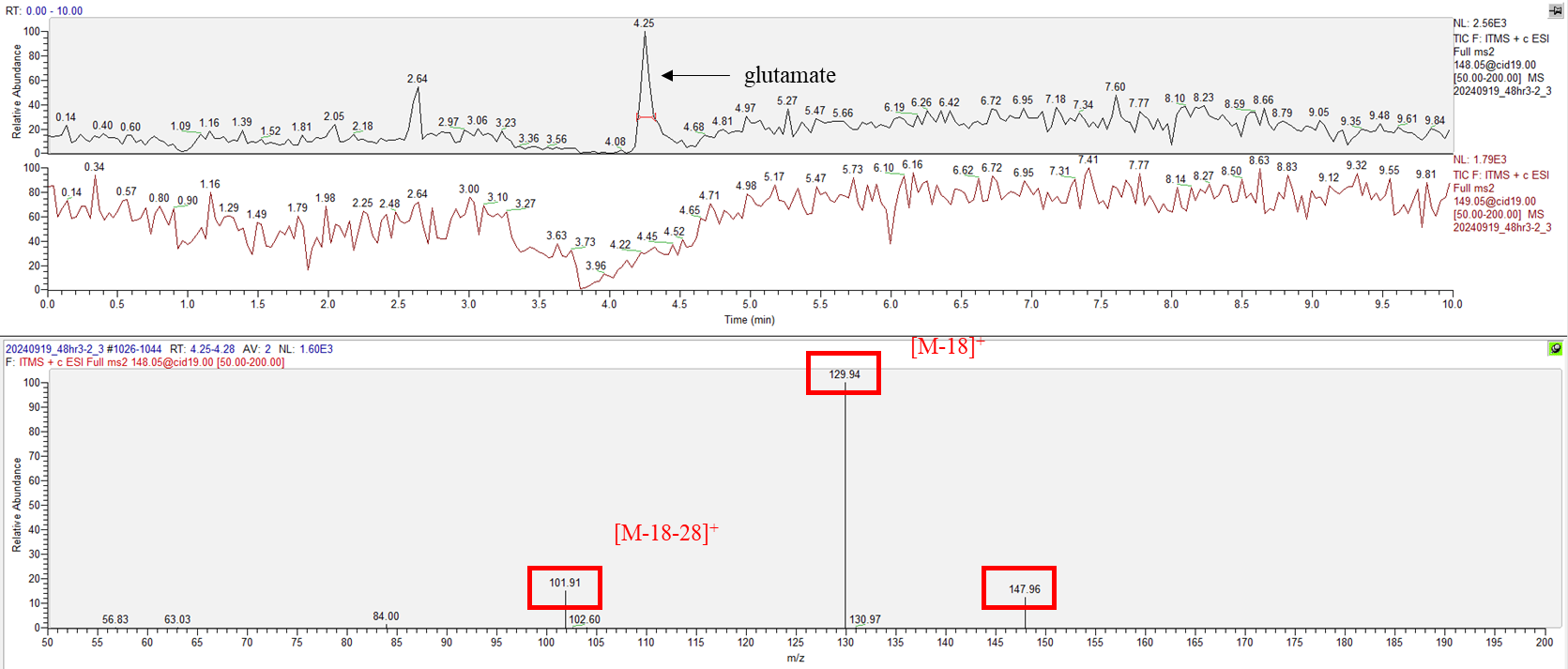

Figure S5. Glutamate synthesis from α-ketoglutarate, sodium dithionite as electron donor. HPLC–MS/MS spectra of reaction mixtures containing α-ketoglutarate, hydroxylamine and sodium dithionite, catalyzed by Fe(0), Fe₃O₄, and artificial [4Fe–4S] clusters with 1M NaCl. The reactions were purged with ^13^CO₂, then incubated at 70 °C for 48 hr.

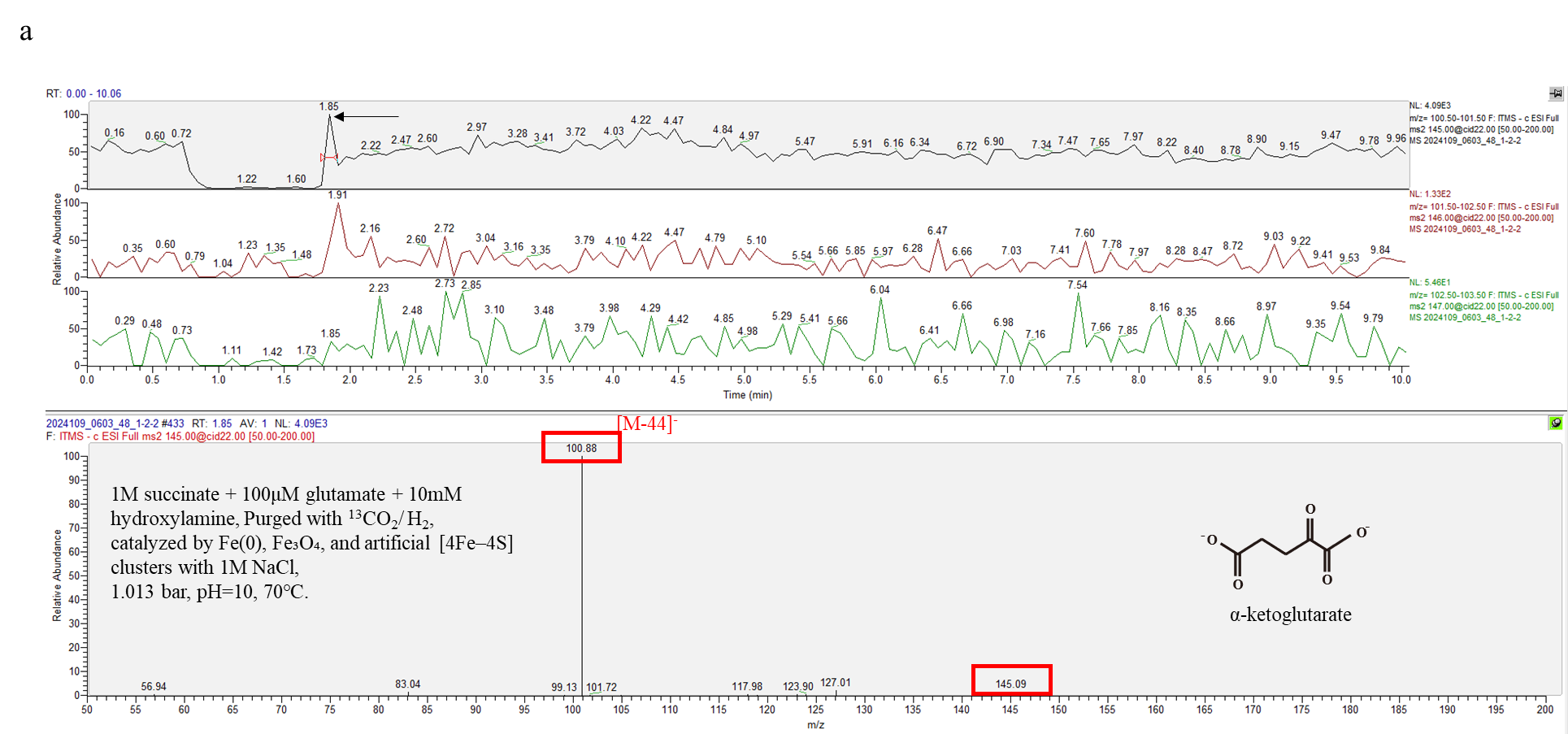

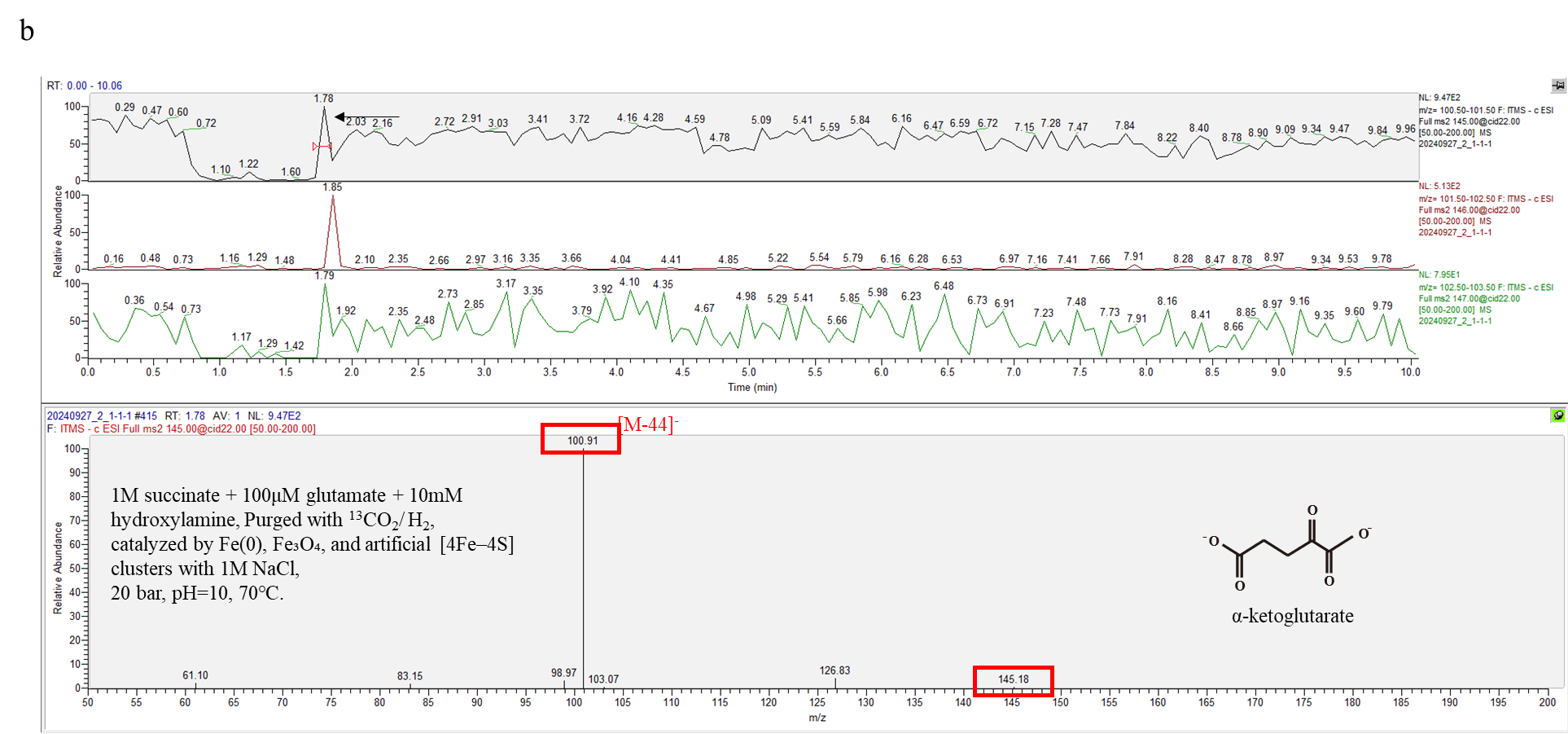

Figure S6. HPLC–MS/MS spectra of m/z 145, the product reaction mixtures containing 1M succinate and 100μM glutamate, 10mM hydroxylamine, catalyzed by Fe(0), Fe₃O₄, and artificial [4Fe–4S] clusters with 1M NaCl. The reactions were purged with ^13^CO₂ and H₂ (2:8), then incubated at 70 °C for 48 hr. (a) the reaction pressure was under 1.013 bar. (b) The reaction pressure was under 20 bars.

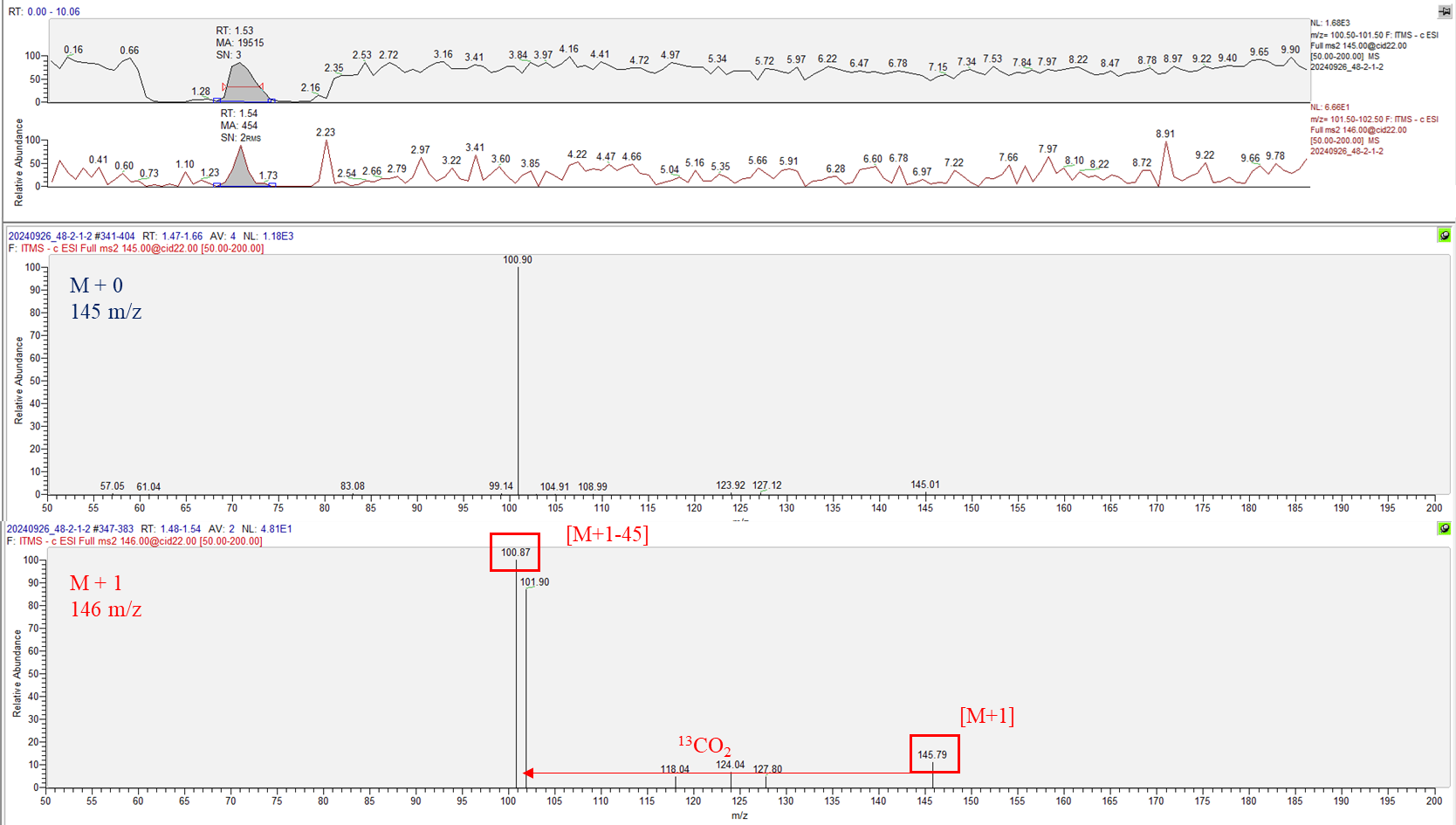

Figure S7. HPLC–MS/MS spectra of m/z 145, and 146. Ion product scan at m/z 100.5~101.5 and m/z 101.5~102.5 from precursor ion m/z 145 (blue), and 146 (red), respectively. The reaction mixtures containing 1M succinate and 100μM glutamate, 10mM hydroxylamine, 10mM sodium dithionite catalyzed by Fe(0), Fe₃O₄, and artificial [4Fe–4S] clusters with 1M NaCl. The reactions were purged with ^13^CO₂, then incubated at 70 °C for 48 hr.

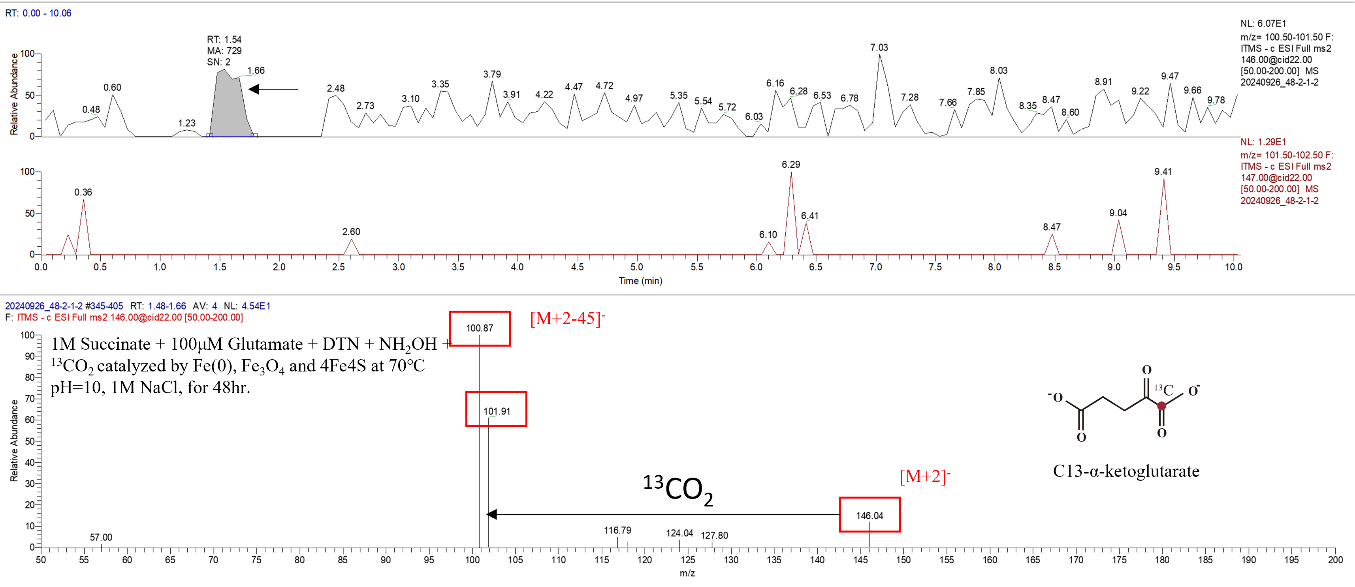

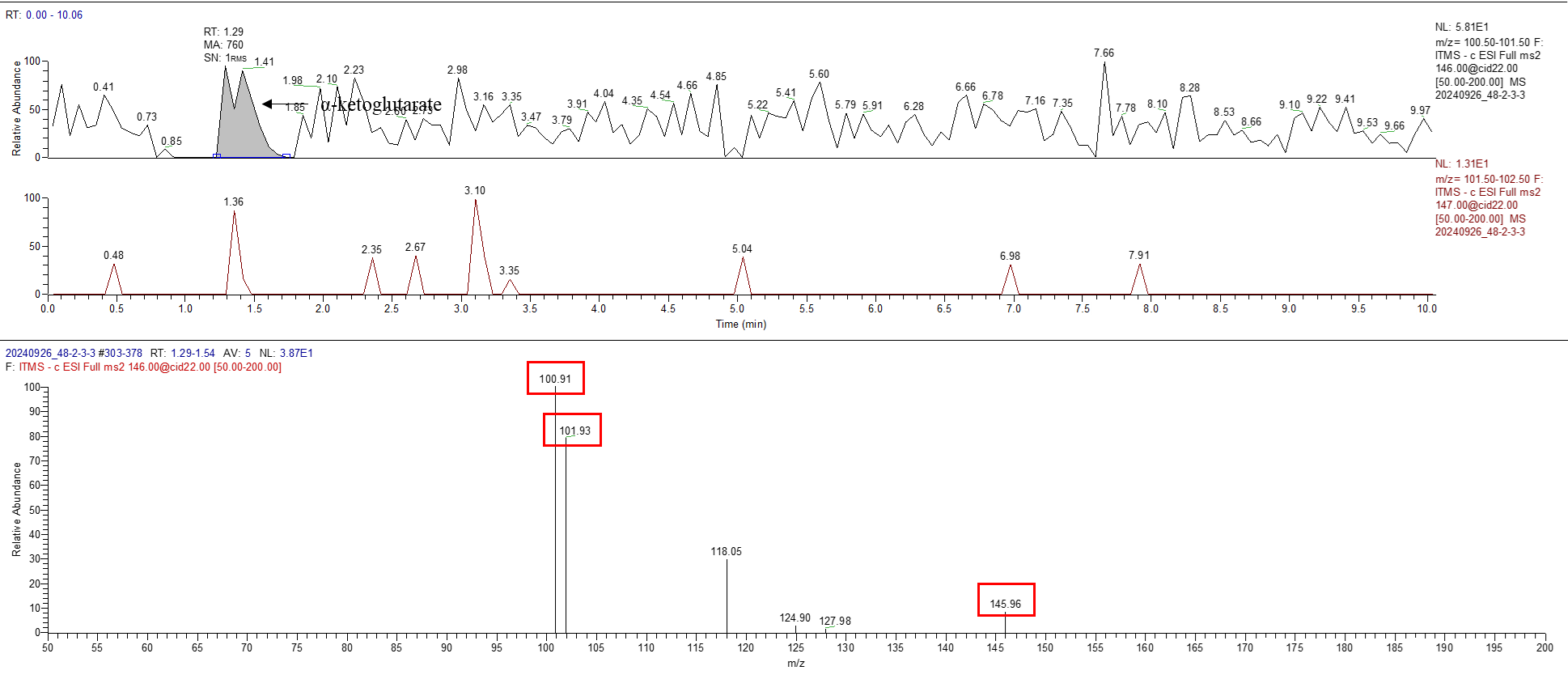

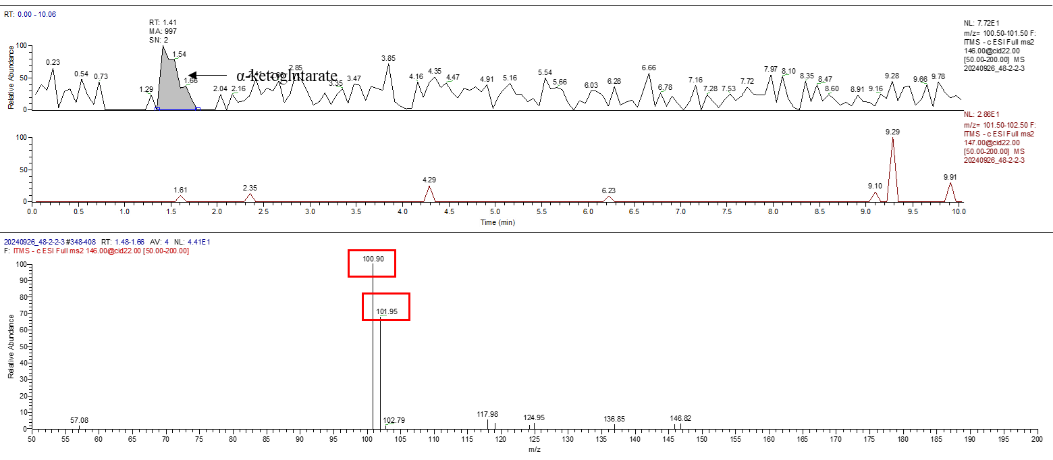

Figure S8. HPLC–MS/MS spectra of m/z 146 to 101. Ion product scan at m/z 100.5~101.5 from precursor ion m/z 146. The reaction mixtures containing 1M succinate and 100μM glutamate, 10mM hydroxylamine, 10mM sodium dithionite catalyzed by Fe(0), Fe₃O₄, and artificial [4Fe–4S] clusters with 1M NaCl. The reactions were purged with ^13^CO₂, then incubated at 70 °C for 48 hr.

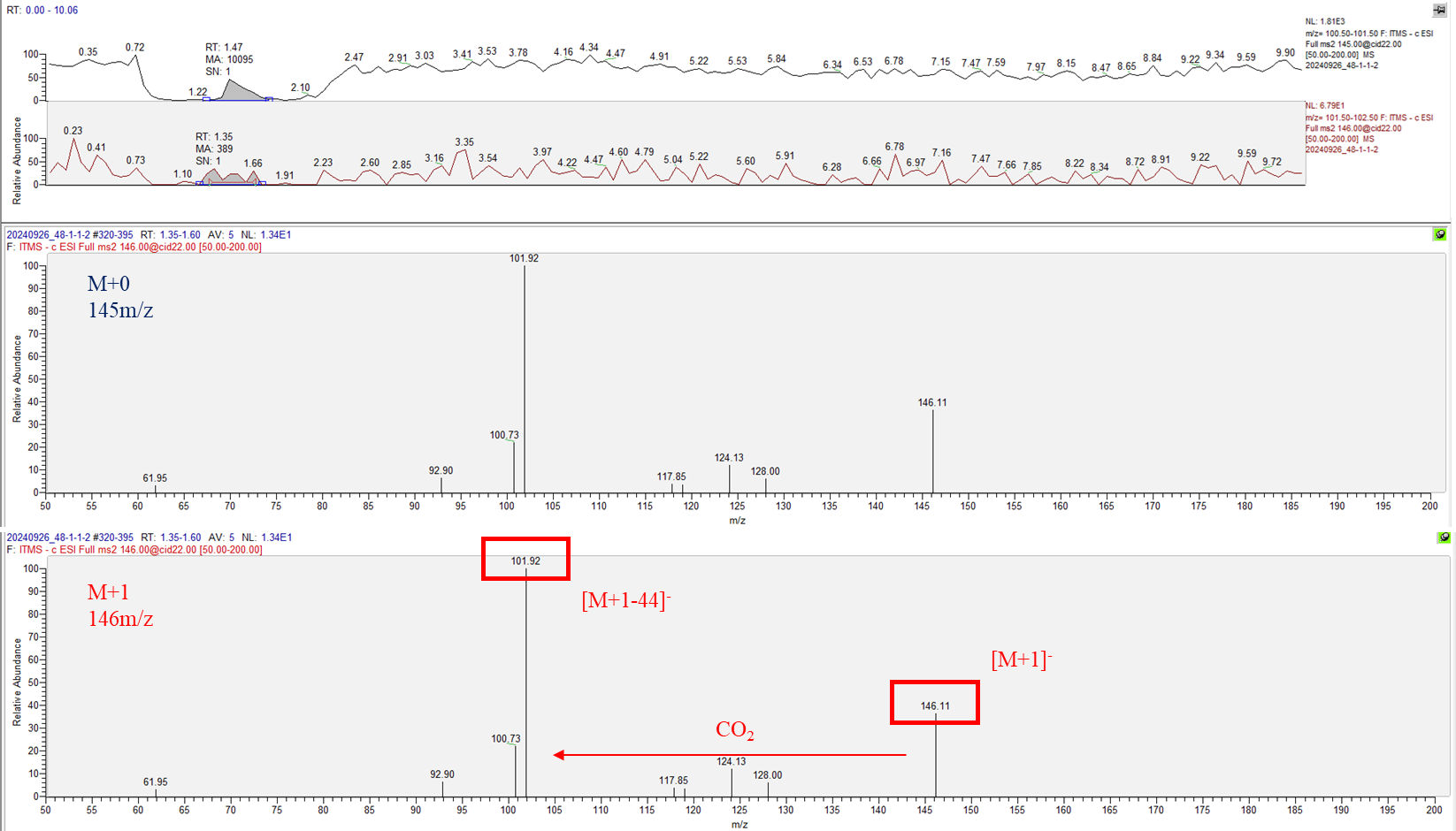

Figure S9. HPLC–MS/MS spectra of m/z 145, and 146. Ion product scan at m/z 100.5~101.5 and m/z 101.5~102.5 from precursor ion m/z 145 (blue), and 146 (red), respectively. The reaction mixtures containing 1M succinate and 100μM glutamate, 10mM hydroxylamine, 10mM sodium dithionite catalyzed by Fe(0), Fe₃O₄, and artificial [4Fe–4S] clusters with 1M NaCl. The reactions were purged with CO₂, then incubated at 70 °C for 48 hr.

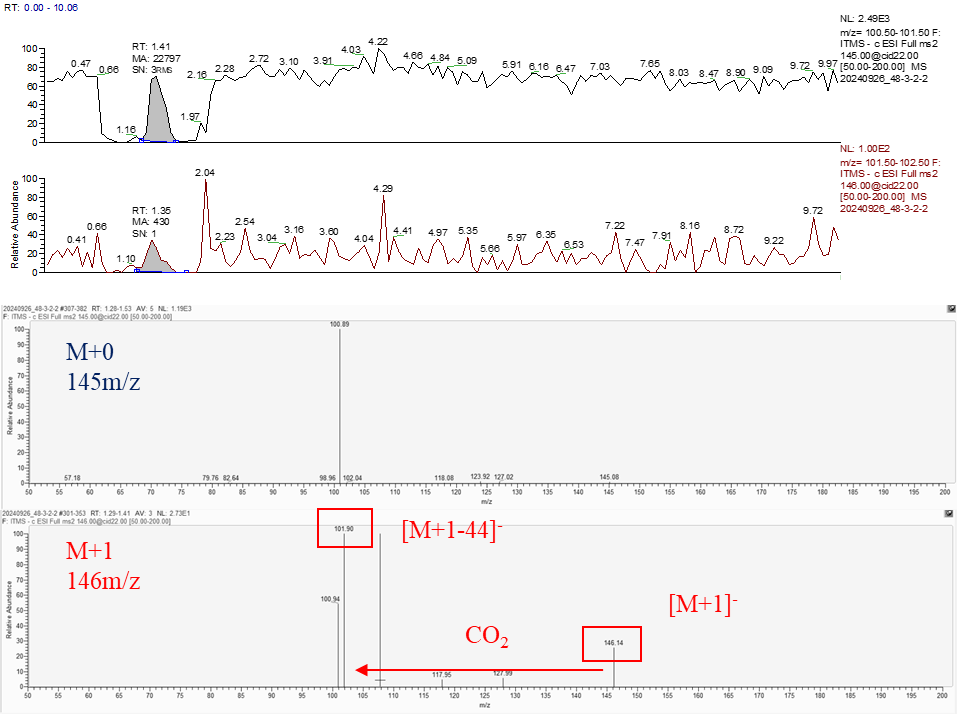

Figure S10. HPLC–MS/MS spectra of m/z 145, and 146. Ion product scan at m/z 100.5~101.5 and m/z 101.5~102.5 from precursor ion m/z 145 (blue), and 146 (red), respectively.The reaction mixtures containing 1M succinate and 100μM glutamate, 10mM hydroxylamine, 10mM sodium dithionite catalyzed by Fe(0), Fe₃O₄, and artificial [4Fe–4S] clusters with 1M NaCl. The reactions were purged with CO₂, then incubated at 70 °C for 48 hr.

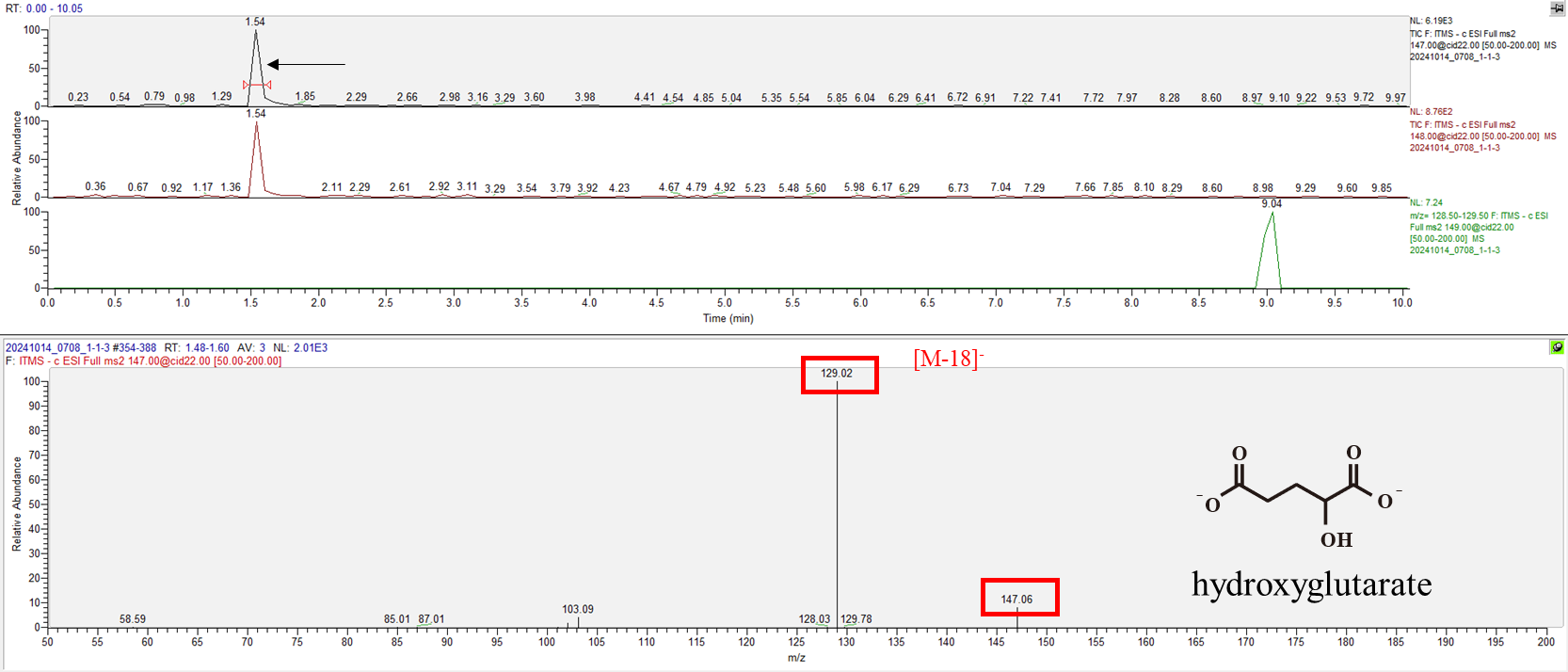

Figure S11. HPLC–MS/MS spectra of m/z 147. The product reaction mixtures contain 10mM succinate and 10mM glutamate, 10mM hydroxylamine, 10mM sodium dithionite catalyzed by Fe(0), Fe₃O₄, and artificial [4Fe–4S] clusters with 1M NaCl. The reactions were purged with ^13^CO₂, then incubated at 70 °C for 48 hr.

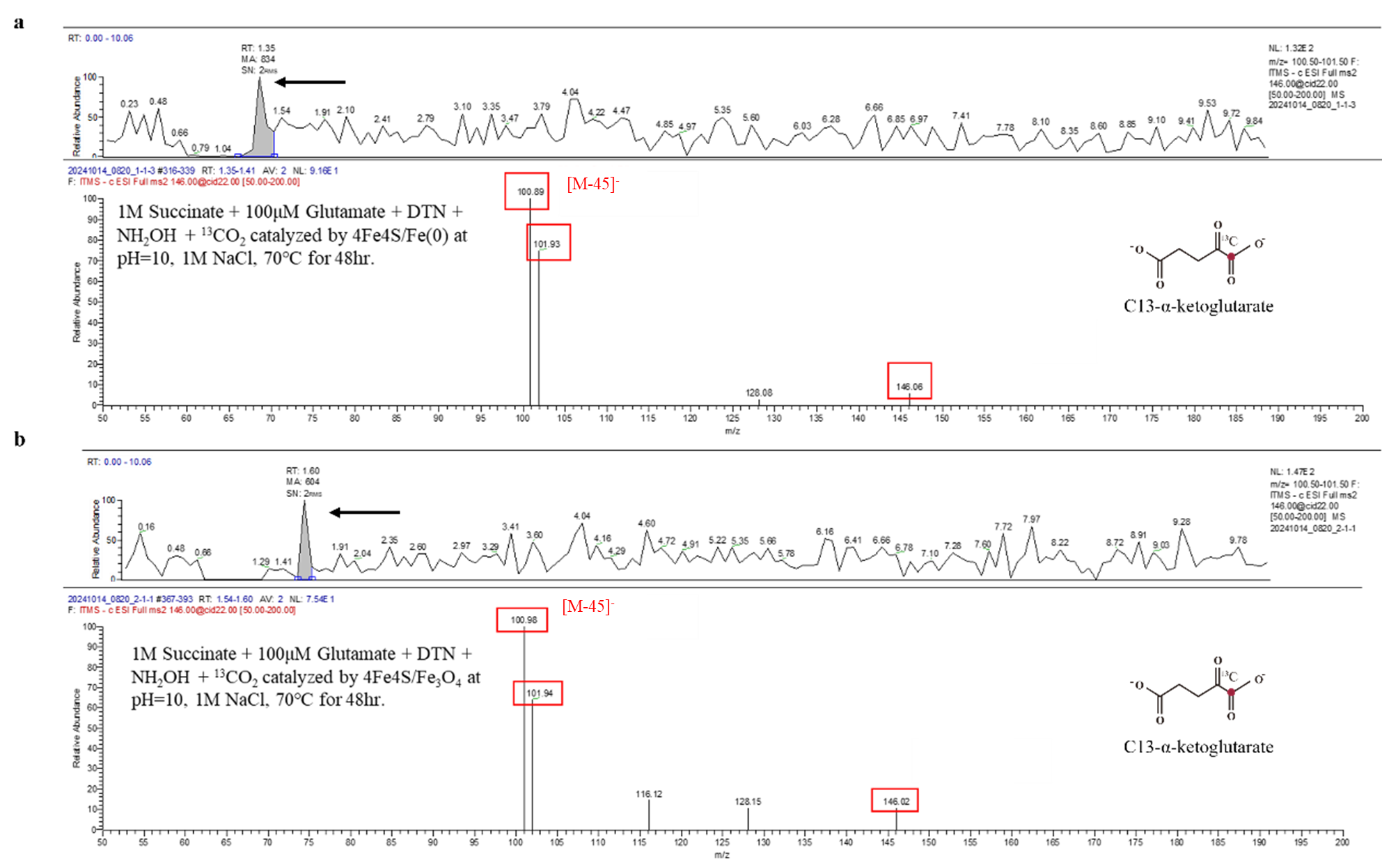

Figure S12. Metal selection reaction. HPLC–MS/MS spectra of m/z 146 to 101, the product reaction mixtures containing 1M succinate and 100μM glutamate, 10mM hydroxylamine, 10mM sodium dithionite catalyzed by (a) artificial [4Fe–4S] clusters and Fe(0) (b) artificial [4Fe–4S] clusters and Fe₃O₄, with 1M NaCl. The reactions were purged with ^13^CO₂, then incubated at 70 °C for 48 hr.

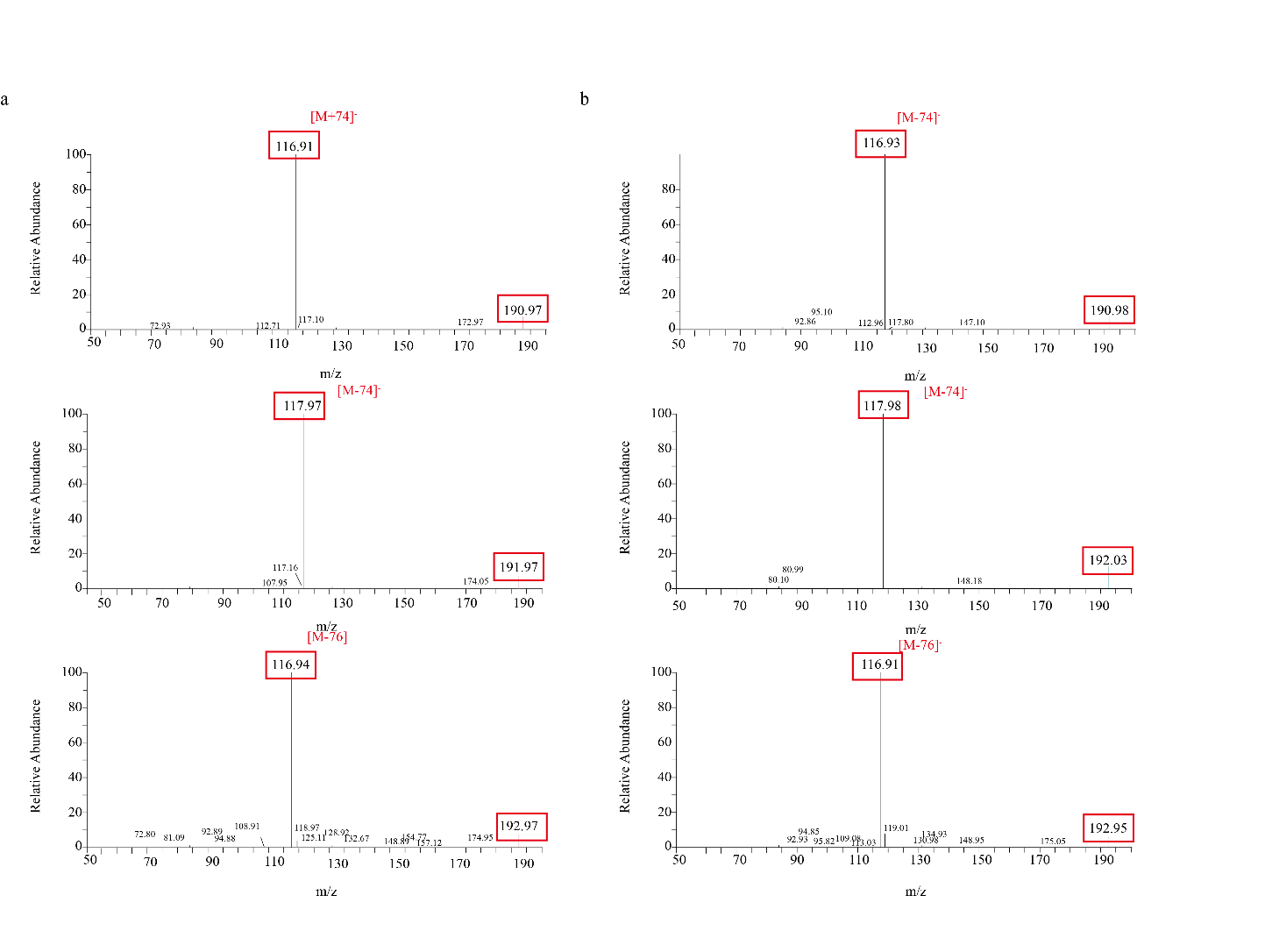

Figure S13. HPLC–MS/MS spectra of m/z 191~193. The product reaction mixtures containing 1M succinate and 100μM glutamate, 10mM hydroxylamine, 10mM sodium dithionite catalyzed by artificial [4Fe–4S] clusters, Fe(0) and Fe₃O₄, with 1M NaCl. The reactions were purged with (a) CO_2_ and (b) ^13^CO₂, then incubated at 70 °C for 48 hr.

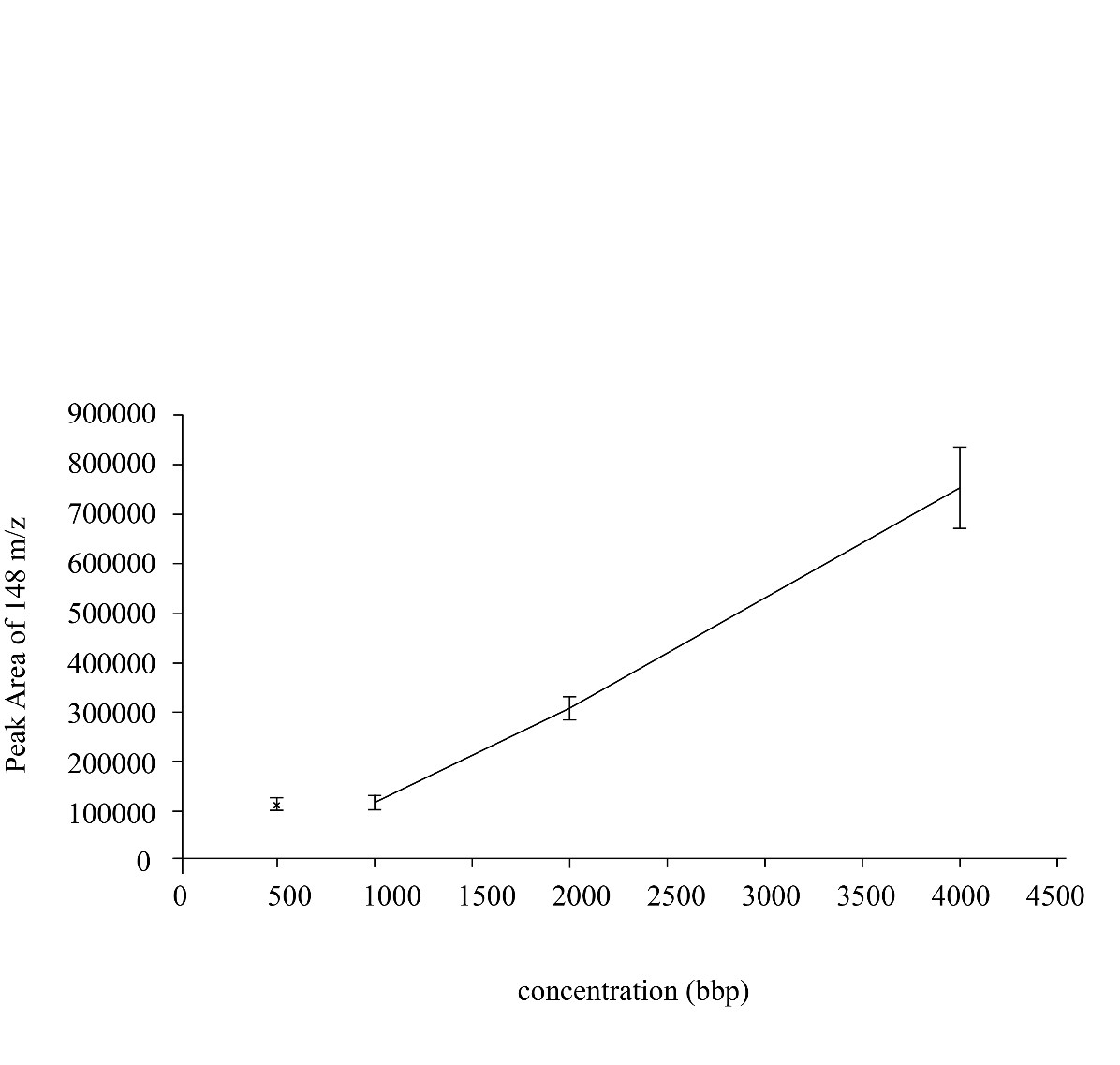

Figure S14. The glutamate standard concentration curve using HPLC-MS/MS for analysis.
