## Supplementary Table for "Nonequilibrium States Promote One-Pot Nonenzymatic Carbon Fixation in the Reverse Tricarboxylic Acid Cycle and Amino Acid Synthesis"

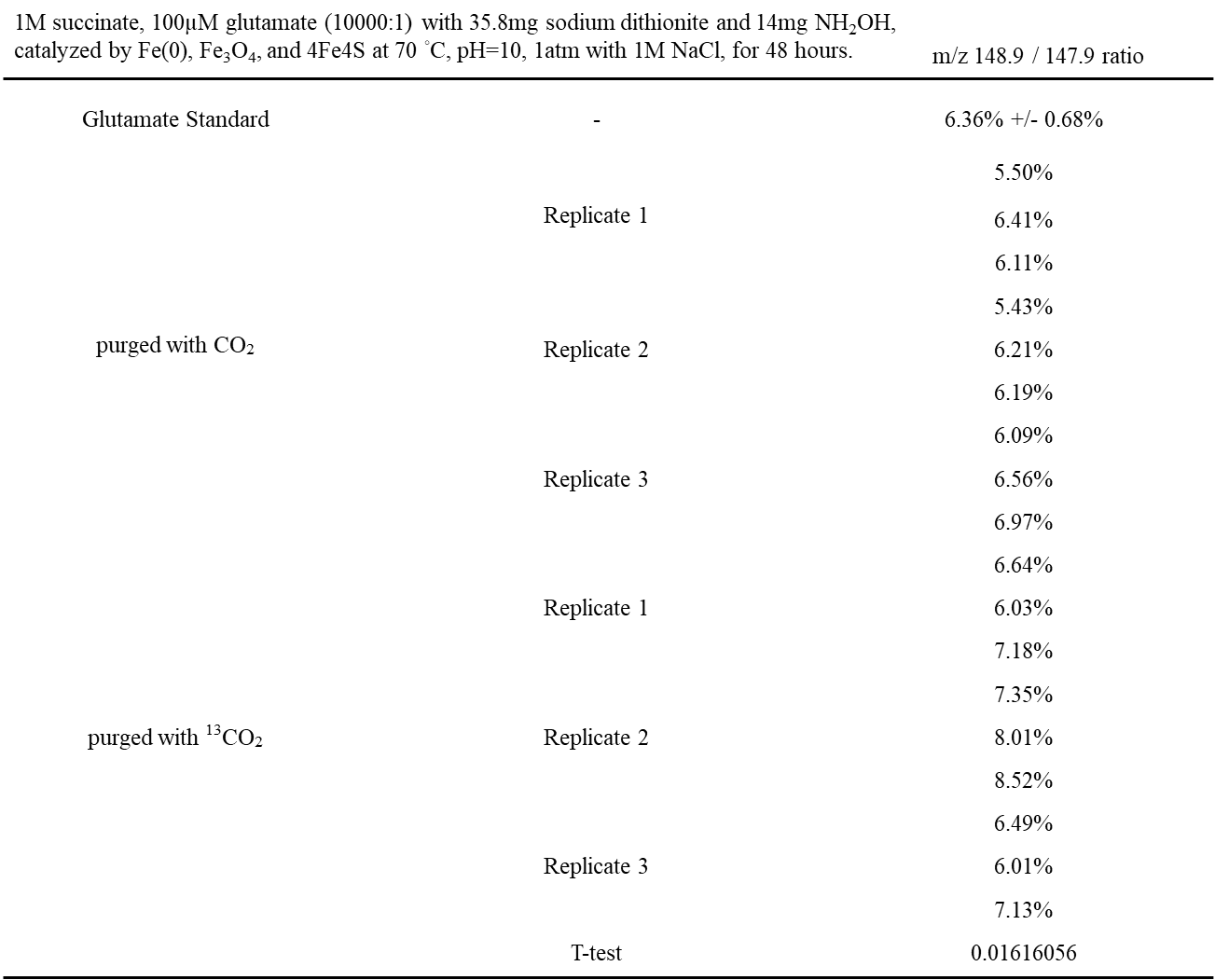
Supplementary Table 1. The m/z 149 to 148 ratio detected through HPLC-MS/MS, the product samples were conducted through 1M succinate, 100μM glutamate with 35.8g sodium dithionite, 14mg NH_2_OH catalyzed by Fe(0), Fe_3_O_4_ and artificial [4Fe4S] at 70℃, pH=10, 1atm with 1M NaCl, then purged with CO_2_ and ^13^CO_2_, the reaction time were 48hr. The standard was purchased from Sigma-Aldrich, >=99%.

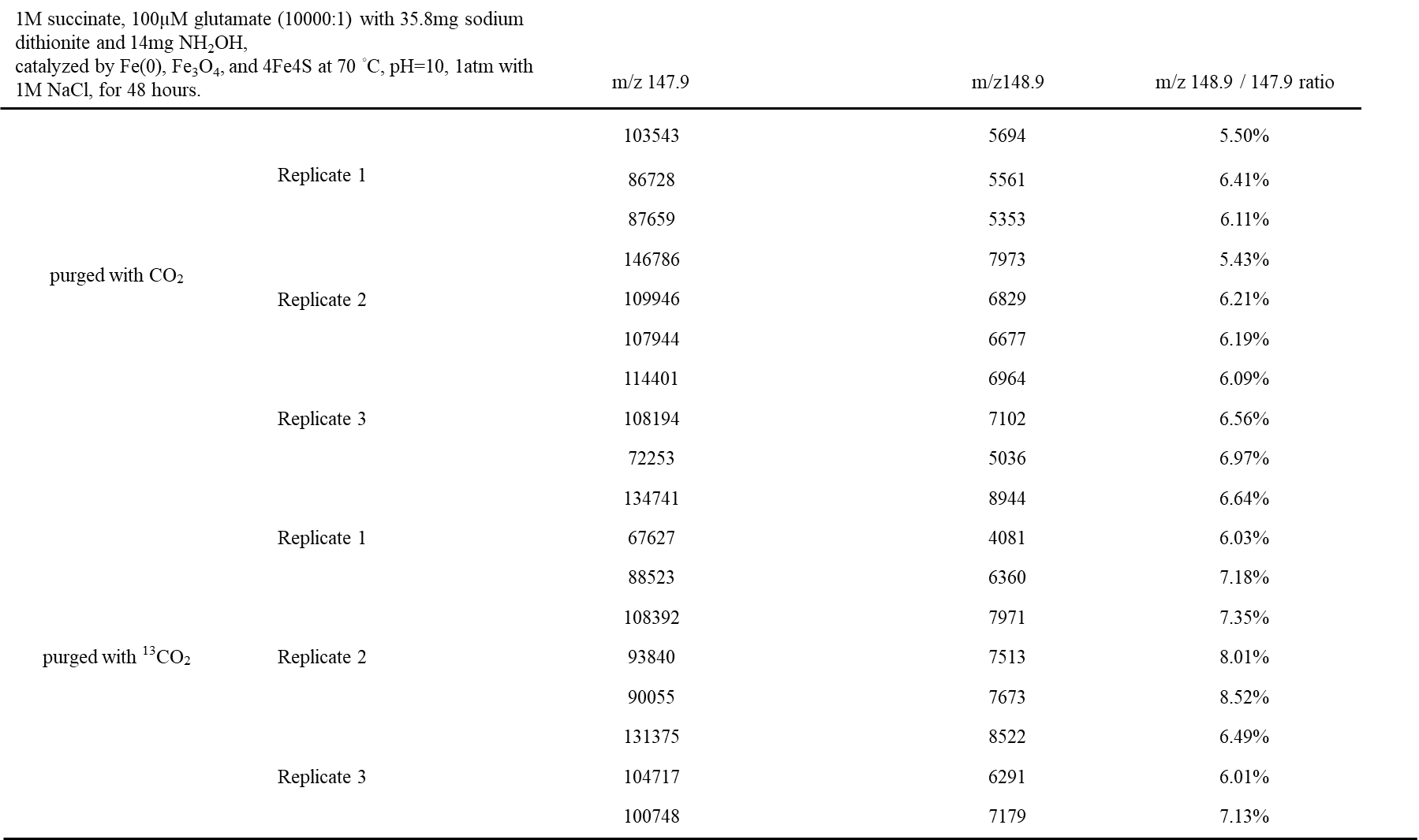

Supplementary Table 2. The peak area of m/z 148.9 and 147.9 detected through HPLC-MS/MS, the product samples were conducted through 1M succinate, 100μM glutamate with 35.8g sodium dithionite, 14mg NH_2_OH catalyzed by Fe(0), Fe_3_O_4_ and artificial [4Fe4S] at 70℃, pH=10, 1atm with 1M NaCl, then purged with CO_2_ and ^13^CO_2_, the reaction time were 48hr. The standard was purchased from Sigma-Aldrich, >=99%.

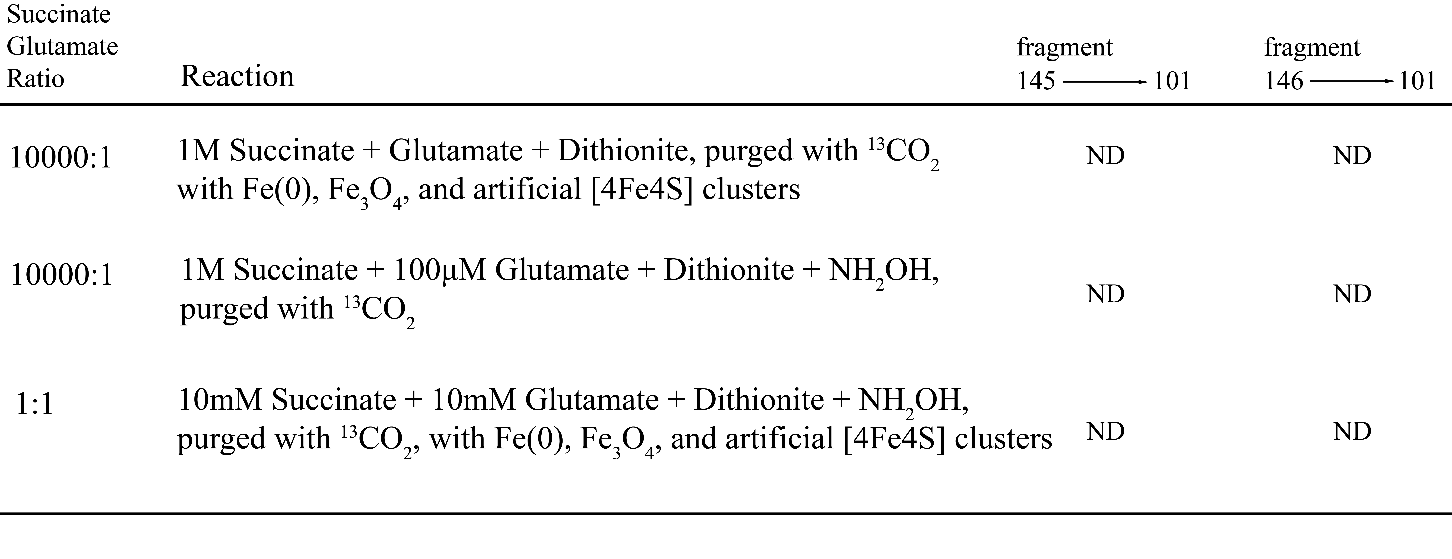

Supplementary Table 3. The results of detecting 145 to 101 m/z and 146 to 101 m/z through HPLC/MS-MS of control group reaction.

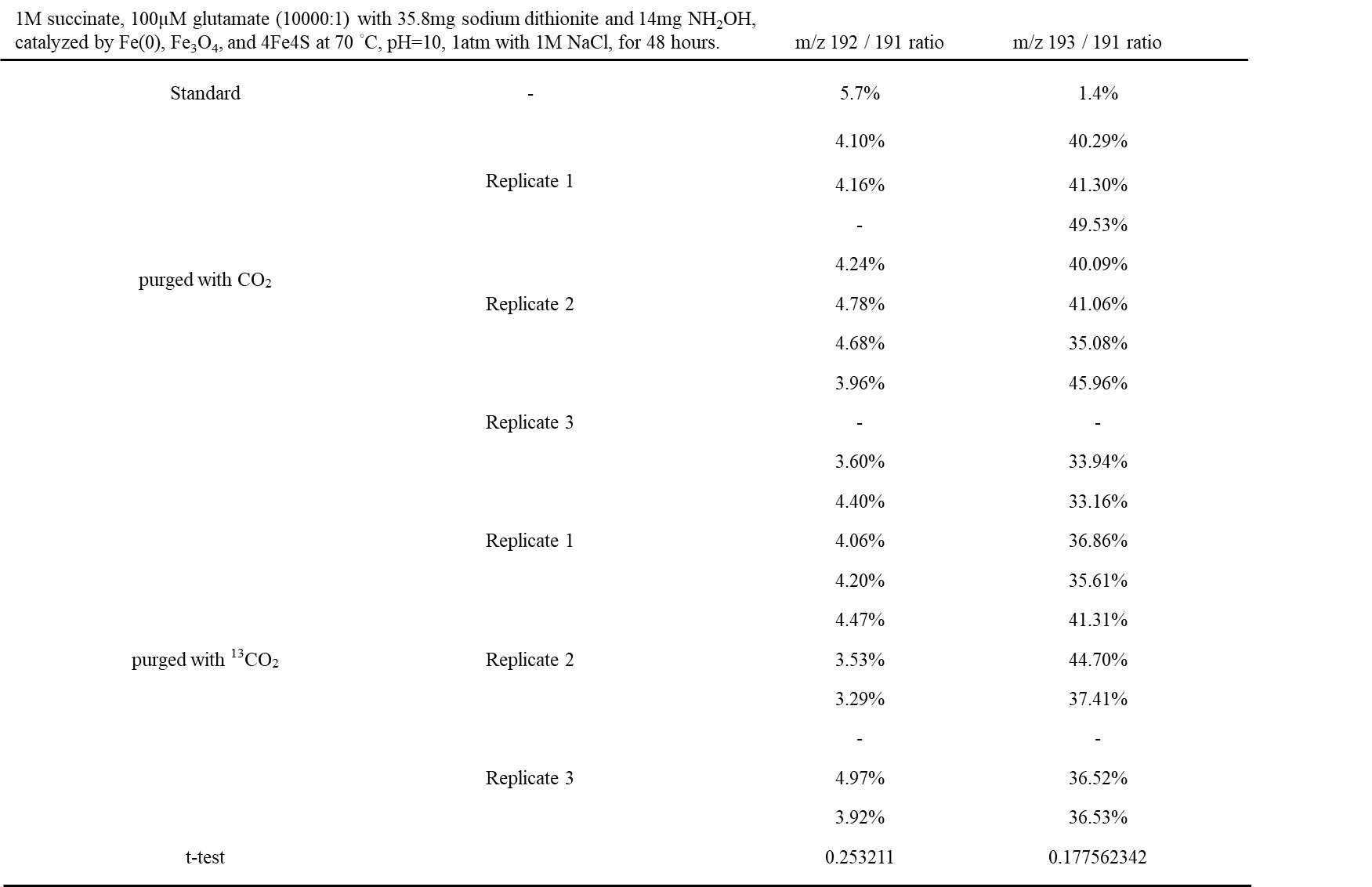

Supplementary Table 4. The m/z 192 to 191 ratio and 193 to 191 ratio detected through HPLC-MS/MS, the product samples were conducted through 1M succinate, 100μM glutamate with 35.8g sodium dithionite, 14mg NH_2_OH catalyzed by Fe(0), Fe_3_O_4_ and artificial [4Fe4S] at 70℃, pH=10, 1atm with 1M NaCl, then purged with CO_2_ and ^13^CO_2_, the reaction time were 48hr.

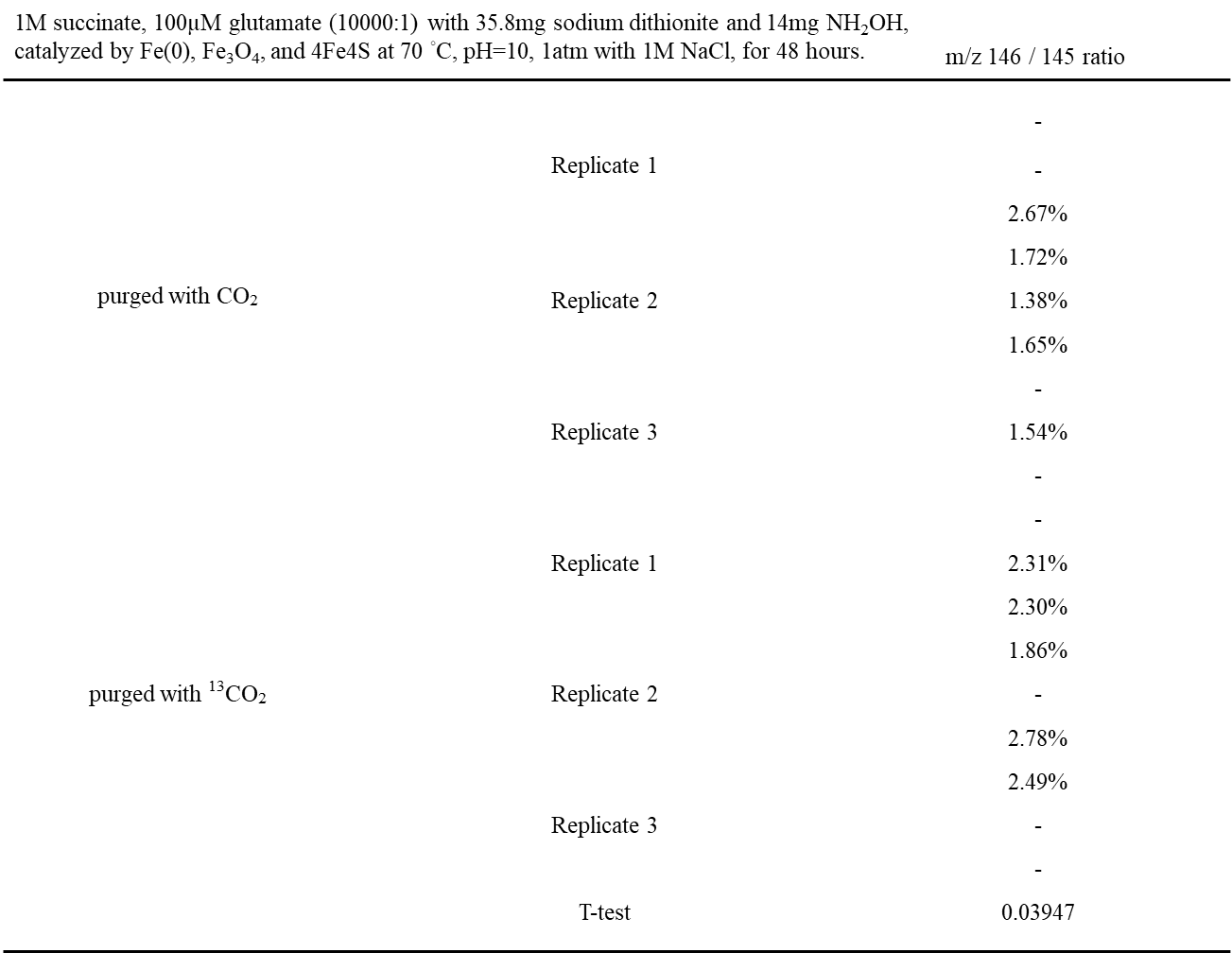

Supplementary Table 5. The m/z 146 and 145 peak area ratio detected through HPLC-MS/MS, the product samples were conducted through 1M succinate, 100μM glutamate with 35.8g sodium dithionite, 14mg NH_2_OH catalyzed by Fe(0), Fe_3_O_4_ and artificial [4Fe4S] at 70℃, pH=10, 1atm with 1M NaCl, then purged with CO_2_ and ^13^CO_2_, the reaction time were 48hr.
